# Self-adjusting Engineered Probiotic for Targeted Tumor Colonization and Local Therapeutics Delivery

**DOI:** 10.1101/2024.01.25.577176

**Authors:** Zhen-Ping Zou, Xin-Ge Wang, Shu-Ting Sun, Jing Mi, Xiao-Peng Zhang, Bin-Chen Yin, Ying Zhou, Bang-Ce Ye

## Abstract

Engineered bacteria have demonstrated great potential for treating a broad array of tumors. However, the precision and safety of controlling the performance of engineered bacteria in vivo remains a central challenge. Here, we utilized genetic circuit programming strategy to construct an engineered *Escherichia coli* Nissle 1917 with accurate targeted colonizing and on-demand payloads releasing ability. The engineered probiotic survives only in the presence of more than 5 mM L-lactate by employing an improved lactate-sensing system, which leads to preventing the growth outside the permissive environments in mice. Meanwhile we introduce an expressing α-hemolysin (SAH) circuit based on quorum-sensing system to augment anti-tumor effect. Furthermore, coagulase induced by high-level lactate creates the closure to deprive tumor of nutrients and oxygen and prevents leakage of bacteria and SAH, which enhances the therapeutic effectiveness and biosafety. This self-adjusting living biotherapeutics significantly inhibits tumor proliferation and prolongs the survival time of colorectal tumor-bearing mice. Together, our work takes a step towards safer and more effective application of living bacteria for tumor treatment in practice.

## INTRODUCTION

The treatment of tumors has been a long-standing challenge. Many strategies have been applied in cancer treatment or clinical testing phase, such as radiotherapy^1^, chemotherapy^2^, CAR-T cell therapy^3, 4^ and living bacterial therapy^5–7^. Especially, due to the selective tumor colonization and high programmability, living bacteria therapies are increasingly being sought after by researchers. Many live tumor-targeting microorganisms, such as *bacillus Calmette-Guérin*^8^, *Salmonella typhimurium*^9^ and *Escherichia coli* Nissle 1917 (EcN)^10^ can colonize in tumor tissues and inhibit tumor cells growth to a certain extent. To augment the antitumor efficacy of bacterial therapy, transport and expression of the therapeutic agents such as various cytotoxic agents^11, 12^, angiogenesis inhibitors^13^, compounds^14^, short hairpin RNA^15^ and chemokines^16^ have been harnessed. Jean *et al.* engineered an arabinose-induced *E. coli* strain to express toxic proteins, which could reduce 91% of the 4T1 tumor volume^17^. Tan *et al.* developed a *S. typhimurium* strain expressing cytolysin A to treat pancreatic cancer^18^. Bacteria also could be designed to express and release IFN-γ, CTLA-4 or PD-L1 nanobodies, and achieved satisfactory effects on tumor treatment^19–21^. However, a critical consideration of using such engineered bacteria in vivo is how to keep them enrich and grow only in tumor. Enhancing the tropism of bacteria via synthetic biology-based genetic circuits is an elegant approach to prevent off-target tissue damage. Yu *et al.* coupled *S. typhimurium* growth with hypoxia to control the colonization in tumors^22^. Another work showed that the engineered *S. typhimurium* with both hypoxia and lactate biosensors via an AND gate improved tumor specificity^23^, but the lactate biosensor based on the *E. coli* LldR regulator and P_lldPRD_ are inhibited partly by glucose and anaerobic^24^ which are the two characteristic hallmarks in the tumor microenvironment (TME). In short, there are still existing several neck bottles needing to be addressed for clinical practice, such as the accuracy and efficiency of anti-tumor drug release, the selective tumor colonization ability, and prevention of therapeutic protein leakage through blood vessels.

To meet these simultaneous demands, we developed an engineered EcN for selective colonization and on-demand recombinant drug protein release in tumor, and this intelligent engineered bacterium prevented the toxic protein from leaking into the healthy tissue by inducing tumor thrombus. Notably, we developed a sensing system specifically responding to high concentrations (> 5 mM) of lactate, and it was not restricted by glucose and hypoxia. This sensing system could regulate the expression of the growth essential gene *asd*^22^ and coagulase^25^ gene simultaneously in an *asd* gene deleted EcN strain. The expression of coagulase causes thrombosis and infarction within the tumor, which results in blocking the nutrients and oxygen supply and preventing the therapeutic protein from leaking. This self-adjusting bacterial system preferentially colonizes in tumors, inhibits tumor proliferation, and prolongs the survival time of tumor-bearing mice (Fig. 1). Our study demonstrates that synthetic biology allows us to reprogram bacteria via integrating multiple genetic circuits to recognize a niche where they should grow and therapeutic proteins could be expressed and released in an effective and safe way. In summary, we established a paradigm for the practical application of living engineered bacteria in the treatment of tumors.

**Fig. 1.**
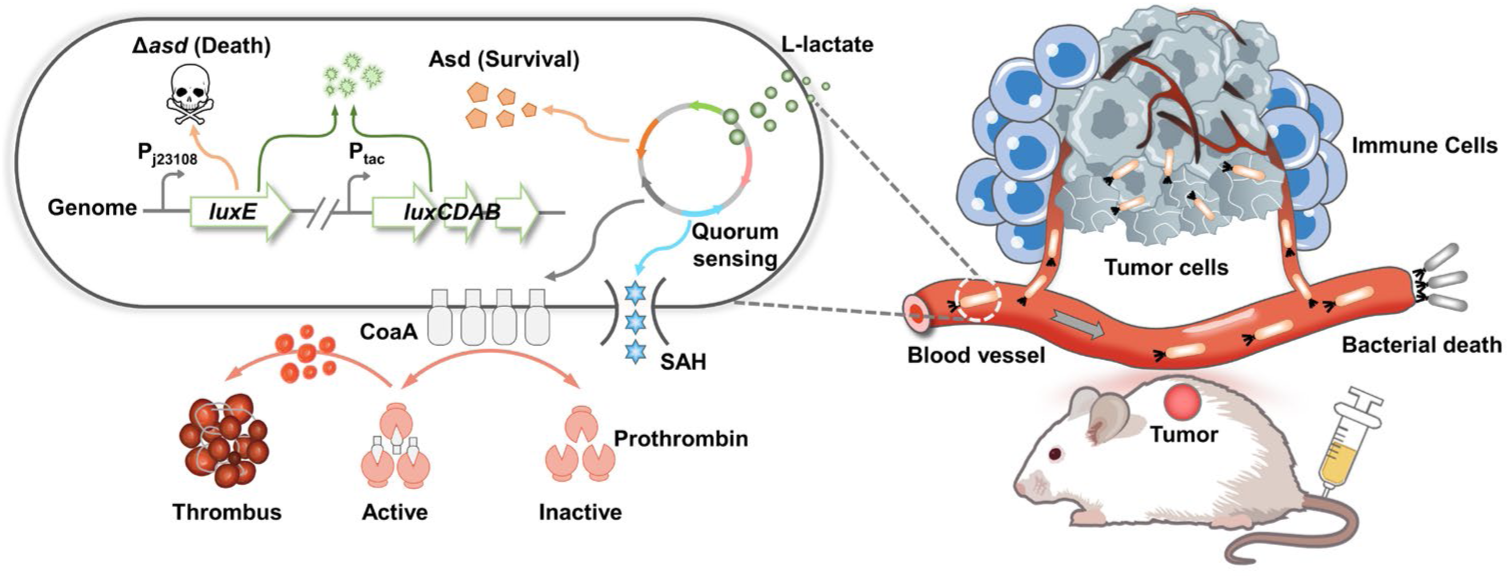
Schematic of self-adjusting engineered EcN for selective tumor colonization and local therapeutics delivery. Left: schematic diagram of the engineered EcN. Right: engineered EcN selectively colonizes tumors and releases payloads via intravenous injection.

## RESULTS

### Engineering Lactate-sensing System to Control Bacterial Growth

Tumor cells uptake glucose and produce lactate at levels typically exceeding 5 mM, in contrast to the normal lactate concentration of approximately 1.5 mM found in healthy organs and blood^26, 27^. We also measured lactate concentrations in diverse tumors, and the results indicated that the lactate contents in tumor is indeed higher than that in normal tissues (Fig. S1 and S2). Therefore, it is feasible to construct an engineered bacterium that strictly relies on high concentration lactate to control tumor targeting and colonization.

We developed lactate sensor by using LldR from *C. glutamicum*, because it is unaffected by glucose and anaerobic^28, 29^. We first optimized the expression of the codon-optimized *lldR* gene using L-arabinose induction system, and use LldR to regulate the reporter sfGFP with various copy numbers and lldRO operator locations to obtain AL, ALO, and ALOO strains (Fig. 2A). The fluorescence intensity decreased or increased with the rising concentrations of L-arabinose and lactate, respectively (Fig. 2B), which indicated that LldR from *C. glutamicum* is a repressor regulated by lactate. Subsequently, we constructed a variants library of biosensors by adjusting the transcription and translation levels of lldR. Most variants exhibited a significant response signal to 5 mM lactate (Fig. 2C). These results indicated that this improved lactate-sensing system could respond to 5 mM lactate and alleviate the inhibitory effect of glucose (Fig. S3A, B).

**Fig. 2.**
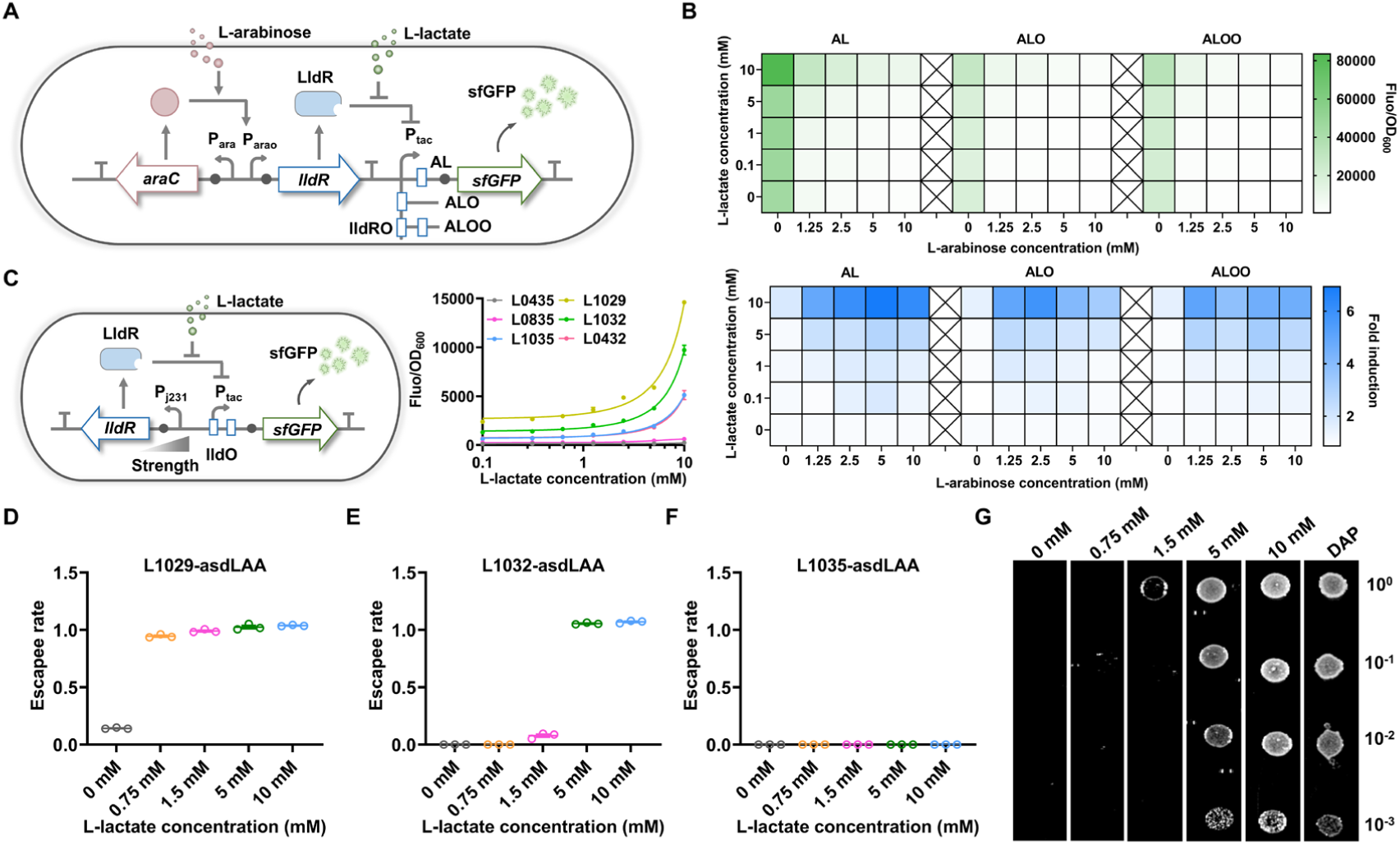
Design and characterization of a lactate inducing system for the bacterial growth. (A) Schematic diagram of tuning the intracellular LldR level by the arabinose inducible system, and the expression of sfGFP is controlled by three promoters, P_AL_, P_ALO_, P_ALOO_ respectively. (B) Fluorescence intensity (top) and the induction fold induced by 0-10 mM arabinose and 0-10 mM lactate after 12 h (bottom). The fluorescence intensity of the culture decreases with increasing arabinose concentration and increases with increasing lactate concentration, indicating that LldR is a repressor. (C) Construction and optimization of a whole-cell biosensor with lactate response. By controlling the transcription and translation levels of LldR, it is convenient to regulate the basal expression and maximum output of the biosensor (mean ± SEM, n = 3). (D-F) Coupling a lactate biosensor with bacterial growth via the expression of an essential gene *asd*. The escapee rate defined as the ratio between colonies grown in DAP and different concentrations of lactate. All strains were cultured for 8 h in lysogeny broth (LB) medium and then plated on LB agar plates with 100 μg/mL DAP (mean ± SEM, n = 3). (G) EcNlA strain at serial dilutions under increasing Lactate levels were cultured for 12 h and observe the bacterial growth.

To effectively control the growth of engineered bacteria under specific conditions, dominating the essential gene expression is an effective strategy. We chose the essential gene (*asd*) under the control of this lactate inducing system. We replaced *asd* in the genome of EcNl with the *luxE* gene to obtain a mutant EcNlE (Fig. S4A, B), and EcNl contained *luxABCD* gene in its genome (refer to our previous work)^30^. EcNlE could grow (Fig. S4C) and produce bioluminescence (Fig. S4D) only when 2,6-diaminopimelic acid (DAP) exceeded 50 μg/mL. In order to utilize lactate to control bacterial growth (Fig. S4E), we tried to place the *asd* gene under the control of three different lactate sensing systems to generate L1029asd, L1032asd and L1035asd strains separately. They all showed approximately 100% escapee rate regardless of lactate from 0 to 10 mM (Fig. S5A). To enhance the escapee rate of engineered bacteria under the limited concentration of lactate, we further reduced the basal expression level of Asd protein by interlinking degradation tags Ssr (AAV or LAA). The results showed that only strain L1032asdLAA (EcNlEA) could grow when the concentration of lactate was more than 5 mM (Fig. 2D-F, Fig. S5B), and the agar plate experiments also confirmed this result (Fig. 2G). Taken together, we successfully developed a lactate-sensing system for precise control of bacterial growth when the lactate is over 5 mM.

### Lactate-induced Display of CoaA on EcN for Coagulation

Targeted delivery of coagulating proteins to tumor blood vessels can block the oxygen and nutrients for tumors and leads to their death. Some coagulation strategies have been used for specific induction of thrombosis within tumors^31,32^. We chose CoaA from *Staphylococcus aureus* to achieve coagulation, because it is convenient to be expressed in prokaryotes and will not promote angiogenesis in vivo^25, 33^. We firstly tested the coagulating ability of purified CoaA (Fig. 3A), and it could coagulate blood within 30 minutes in vitro (Fig. 3B). Then, we integrated the *coaA* gene with lactate inducing system to generate L1035coaA strain (Fig. 3C). L1035coaA was induced with different lactate concentrations for 6 h (OD_600_ ≈ 0.4), and only the cultures induced by lactate with 5 and 10 mM could cause blood coagulation in 3 h (Fig. 3D). It was also worth noting that CoaA was embedded on the cell membrane, rather than secreted into the extracellular environment (Fig. 3E).

**Fig. 3.**
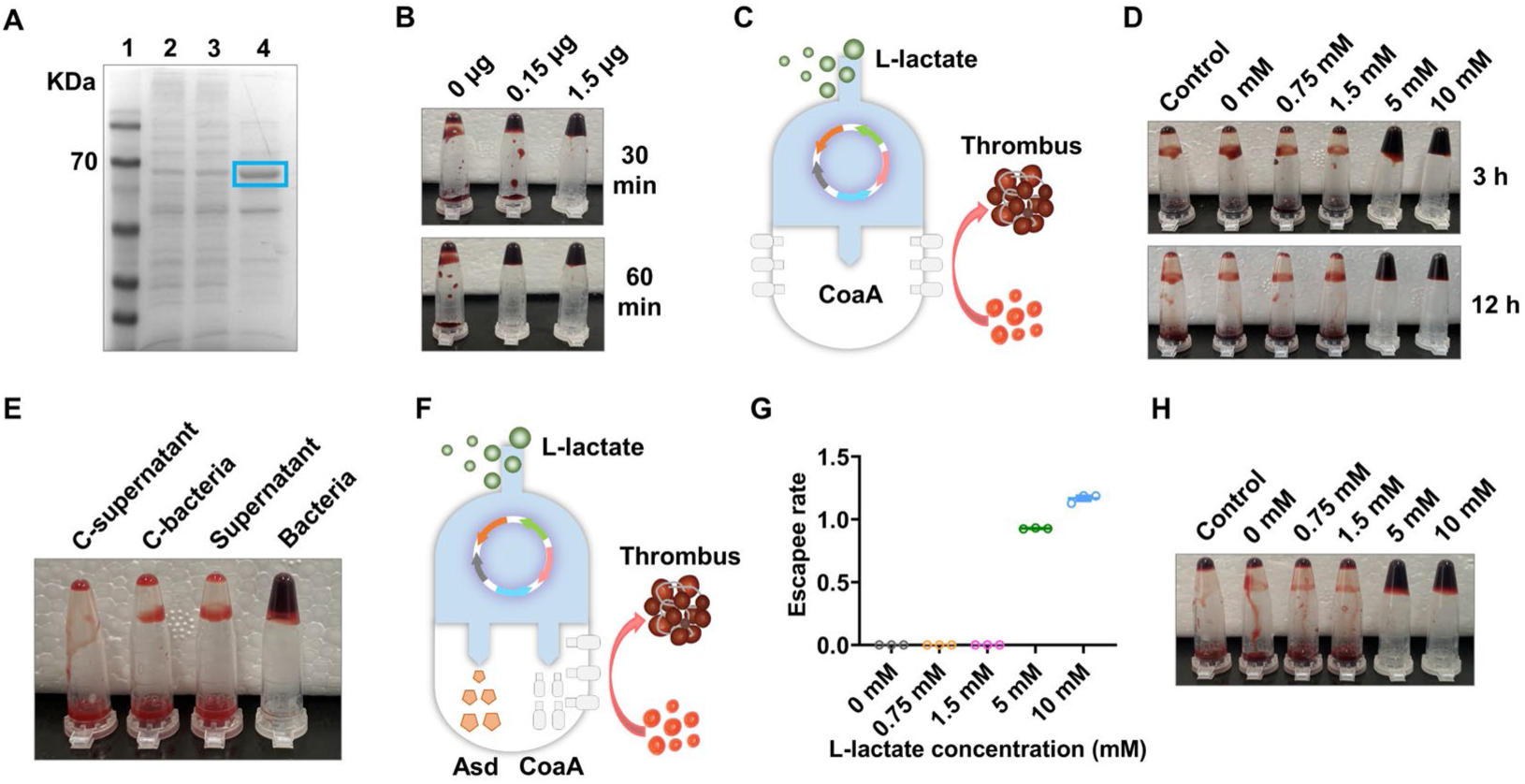
Simultaneous control of bacterial growth and coagulase expression through lactate biosensor. (A) Expression and purification of truncated CoaA coagulase. (B) Images of the coagulation formation at 30 and 60 min after being treated by different concentrations of CoaA. (C) Construction of an engineered EcN strain with lactate induced coagulase expression. (D) lactate responsive engineered EcN L1035coaA induced blood coagulation. Pre-culture L1035coaA strain (with 100 μg/mL DAP) to OD_600_ ≈ 0.4, then induce for 6 h with different concentrations of lactate. Mix 50 μL cultures with 50 μL blood and incubated at 37°C for 3 and 12 h to test blood coagulation. (E) Determine the type of coagulase. Induce pre-cultured L1035coaA with 0 mM or 10 mM for 6 h, centrifuge to collect culture medium supernatant and bacterial precipitation. Mix 50 μL supernatant and bacteria (resuspended with 50 μL PBS) with 50 μL blood and incubate at 37°C for 12 h, then observe the blood clotting. (F) Construction of an engineered EcN strain with lactate induced bacterial growth and coagulase expression. (G) The growth of EcNlEAC strain is strictly regulated by lactate concentration. EcNlEAC strain can growth only in permissive lactate levels (more than 5 mM) (mean ± SEM, n = 3). (H) The expression of coagulase in EcNlEAC strain also strictly regulated by lactate concentration.

In order to construct a strain capable of controlling growth and expressing coagulase induced by lactate simultaneously, we integrated *lldR*, *asd-laa* gene and *coaA* gene into one plasmid to generate an engineered bacterial library (Fig. 3F). Ultimately, we obtained a strain named EcNlEAC that could selectively grow (Fig. 3G) and coagulate blood (Fig. 3H) under permissive conditions (lactate exceeds 5 mM). Please refer to Table S1 for the characteristics of EcNlEAC.

### Cytotoxic Protein Enhances Anti-tumor Effect

To further improve the anti-tumor effect, we integrated cytotoxic protein SAH with the quorum-induced system into EcNlEAC. Quorum-sensing (QS) bacteria have the ability to modulate specific gene expression in response to the population density^34^. This feature endows our engineered bacteria with ability to express and release payloads only in tumor. We first demonstrated the effectiveness of QS system through fluorescent proteins (Fig. S6). Then we constructed the expressing SAH^17^ by QS system in EcNlEAC to obtain strain named EcNIEACS containing two systems: i) Lactate-sensing system strictly controlling bacterial growth and coagulase expression, and ii) QS system inducing toxic protein SAH expression. (Fig. 4A).

**Fig. 4.**
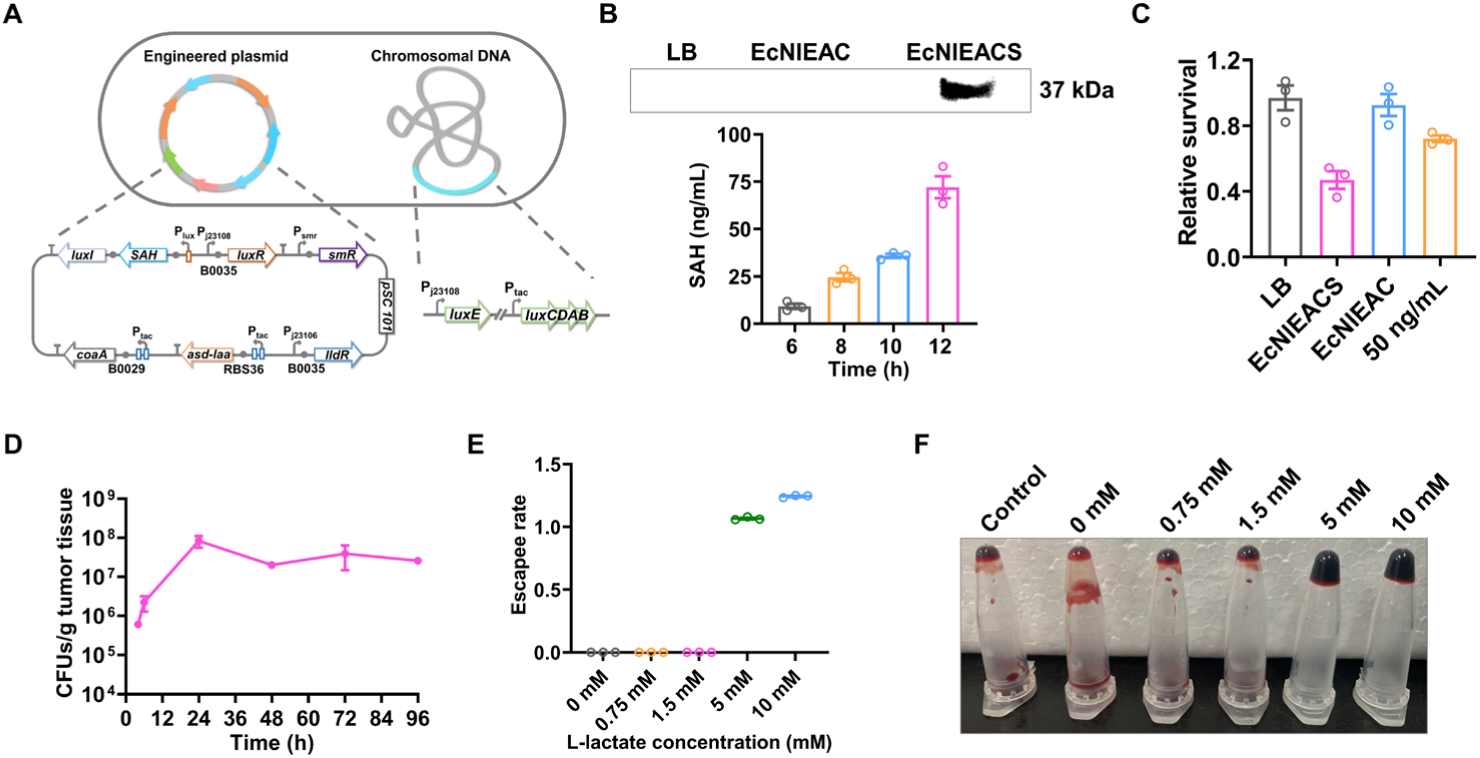
Design and functional characterization of a multifunctional EcN. (A) Genetic circuit diagram of EcNlEACS. (B) Detection and quantification of the secretion level of SAH by WB and ELISA. EcNlEAC and EcNlEACS culture supernatant were collected by centrifugation for WB analysis after 12 h culture, use LB medium as a control. Only the culture supernatant derived from EcNlEACS showed a specific binding band (top). Detection of SAH levels in culture supernatant through ELISA during the cultivation process (bottom) (mean ± SEM, n = 3). (C) Cytotoxicity analysis through CCK-8. Supernatant from EcNlEACS cultures decreased survival of MCF-7 cells compared to LB control, EcNlEAC and 50 ng/mL SAH (mean ± SEM, n = 3). (D) Quantification of EcNlEACS colonization in MC38 tumors at different time points after injection of 1.5 × 10^7^ CFU bacteria (mean ± SEM, n = 3). (E) Escape rate of EcNlEACS strain at various lactate concentrations, EcNlEACS strain can growth only in more than 5 mM lactate levels (mean ± SEM, n = 3). (F) Coagulation ability of EcNlEACS strain induced by various lactate concentrations.

Next, to analyze the secretion and quantitative determination of SAH, western blot (WB) and enzyme-linked immunosorbent assay (ELISA) were carried out. As shown in Fig. 4B, only the medium of EcNlEACS displayed a specific band compared with other groups, and the secretion of SAH could reach at 75 ng/mL after 12 h cultivation. To test the ability of EcNlEACS killing cancer cells, the supernatants of EcNlEAC and EcNlEACS were co-incubated with cancer cells. The supernatant of EcNlEACS reduced cell viability sharply (about 60%) after 6 h incubation (Fig. 4C). Noticeably, the percent of the dead cell in the EcNlEACS group was significantly higher than purified SAH with 50 ng/ml (Fig. 4C). The blood agar test also confirmed the functional SAH produced by EcNlEACS (Fig. S7). Finally, we evaluated the colonization ability of EcNlEACS in tumors. EcNlEACS (1.5 × 10^7^ CFU) was intravenously injected to female C57BL/6 mice bearing subcutaneous MC38 tumors, and sacrificed at 4, 6, 24, 48, 72, 96 h after injection. The tumor tissues were collected, homogenized, and plated on LB plates after serially diluted, there were still 1×10^7^ CFU EcNlEACS in tumors even after 96 h (Fig. 4D). The engineered bacterial growth and the coagulase expression were also investigated at different lactate concentrations from 0 to 10 mM, and both of them were strictly controlled by high concentration lactate (Fig. 4E, F).

### Engineered Probiotic EcNlEACS Exhibits Biosafety

To evaluate the biocontainment and safety of our engineered bacteria, we intravenously injected 1.5 × 10^7^ CFU EcNllux, EcNlEA, EcNlEAC, EcNlEACS to healthy mice and sacrificed at 24 h and 48 h respectively. Wherein, EcNllux harbored a plasmid expressing the *luxE* gene to serve as a negative control. Liver, spleen and lung were collected, homogenized, and plated on LB plates after serially diluted to calculate the number of engineered bacteria. Comparatively, lactate-dependent strains exhibited a 1-2 order decrease in CFU when recovered from the liver and spleen, and a 2-3 order decrease when recovered from the lung, as compared to the control strain EcNllux. The results showed that the recovered CFU of all lactate-dependent strains were much lower than strain EcNllux in the organs (Fig. 5A-C and Fig. S8A-C). Moreover, EcNlEACS group exhibited a slightly weight loss compared to EcNllux after intravenously injection (Fig. 5D).

**Fig. 5.**
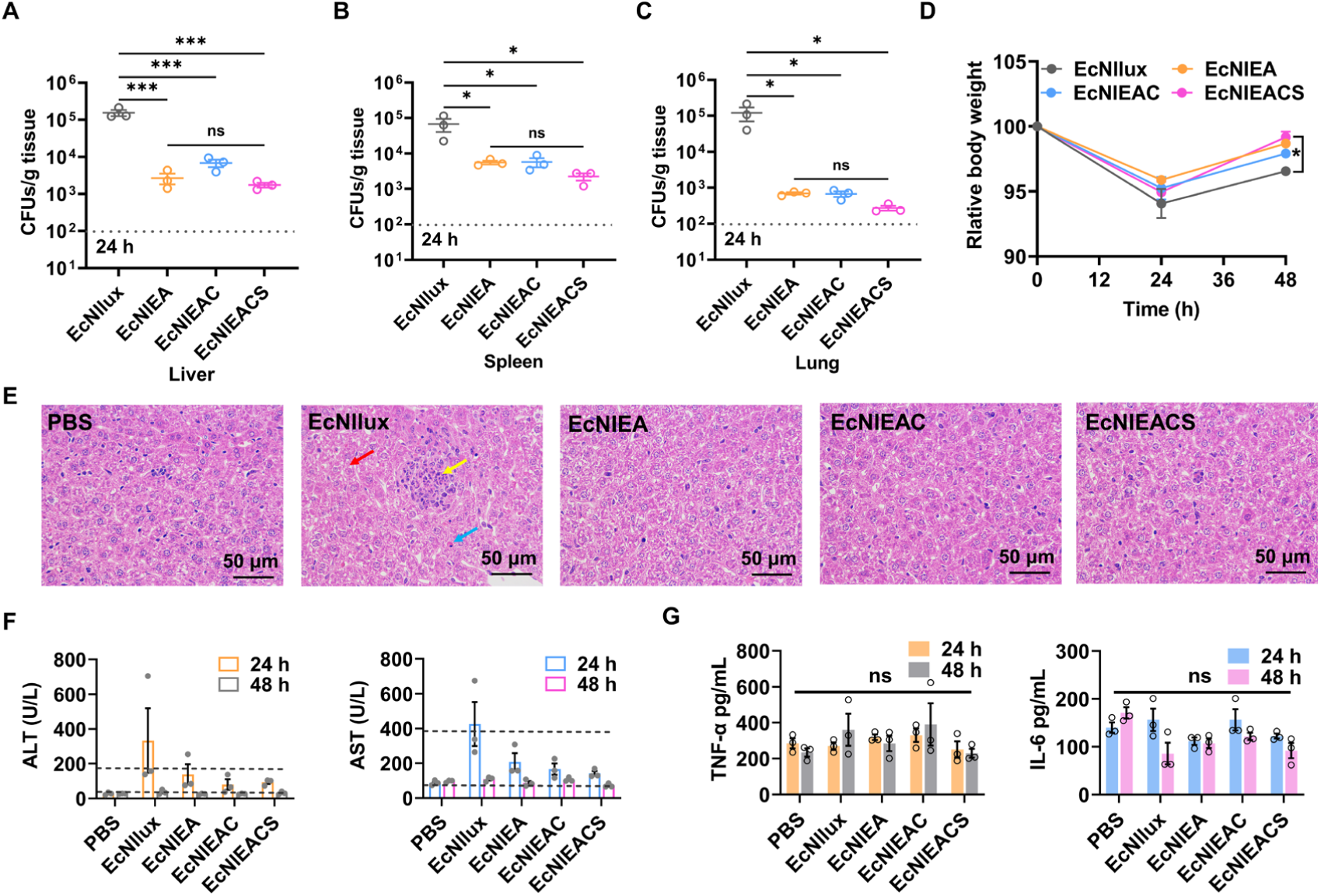
The biocontainment and safety of the engineered EcN. (A-C) Quantification of bacterial colonization in different organs (liver, spleen, lung) harvested from healthy mice after inject bacterial for 24 h. LOD = 1×10^2^ CFU/g (mean ± SEM, n=3). Statistical significance was determined by a one-way ANOVA, *p ≤ 0.05, ***p ≤ 0.001. (D) Weight changes in healthy mice during different strains treatment (mean ± SEM, n=3). Statistical significance was determined by a two-way ANOVA, *p ≤ 0.05. (E) Histological images of liver sections from each group stained with hematoxylin and eosin. Red arrow: mild edema of liver cells. Yellow arrow: hepatic sinus stenosis and small focal necrosis. Blue arrow: hepatocyte eosinophilic transformation. (F) Serum transaminase levels in healthy mice. Detection of transaminase levels in mouse serum after 24 and 48 h of 1.5 × 10^7^ CFU bacterial injection (mean ± SEM, n = 3). (G) Serum inflammatory factors levels in healthy mice. Detection of inflammatory factors levels in mouse serum after 24 and 48 h of 1.5 × 10^7^ CFU bacterial injection via ELISA (mean ± SEM, n = 3). Statistical significance was determined by a one-way ANOVA, ns p > 0.05.

To evaluate the damage of liver, we observed the histopathological status by hematoxylin and eosin (H&E) histological staining after 24 h of intravenous injection of PBS, EcNllux, EcNlEA, EcNlEAC and EcNlEACS. The control group (EcNllux) showed a degree of pathological characteristics, such as mild edema of liver cells (red arrow), hepatic sinus stenosis and small focal necrosis (yellow arrow) and hepatocyte eosinophilic transformation (blue arrow), whereas less pathological characteristics were observed in all lactate-dependent strains groups (Fig. 5E). We also measured the levels of alanine transaminase (ALT), aspartate aminotransferase (AST) and cytokines in the serum. Serum transaminase levels of EcNIlux group reached the upper limit of healthy ones, while other groups were almost within normal range after 24 h administration (Fig. 5F). There was no significant difference in cytokine levels among all groups after 24 h administration (Fig. 5G). Collectively, these results demonstrated that EcNlEACS could reduce tissue off-target and owned satisfied biosafety.

### Distribution of Engineered Probiotic EcNlEACS in Vivo

To evaluate the colonization ability in tumor, EcNlEACS and control strains (1.5 × 10^7^ CFU) were intravenously injected to C57BL/6 mice bearing subcutaneous MC38 tumors. Organs and tumors were homogenized to evaluate bacterial colonization ability for 24 h. All strains colonized tumors at a comparable level (10^7^-10^8^ CFU/g tissue), but the abundance of EcNIEACS in the liver and spleen is two orders of magnitude lower compared with EcNIlux (Fig. 6A-C). We then computed a dose-toxicity curve and demonstrated that lactate-dependent strains have a higher maximum tolerated dose (MTD) than EcNllux in healthy mice (Fig. 6D). Moreover, using imaging of bioluminescent bacterial populations, we could identify bacterial populations in tumors obviously (Fig. 6E). Lactate responding strains at MTD showed rapid weight recovery within three days (Fig. 6F). We next injected EcNllux, EcNlEA, EcNlEAC and EcNlEACS intravenously at the corresponding MTD of each strain (2.9 × 10^7^ CFU, 8.5 × 10^7^ CFU, 8.9 × 10^7^ CFU and 9.7 × 10^7^ CFU for

**Fig. 6.**
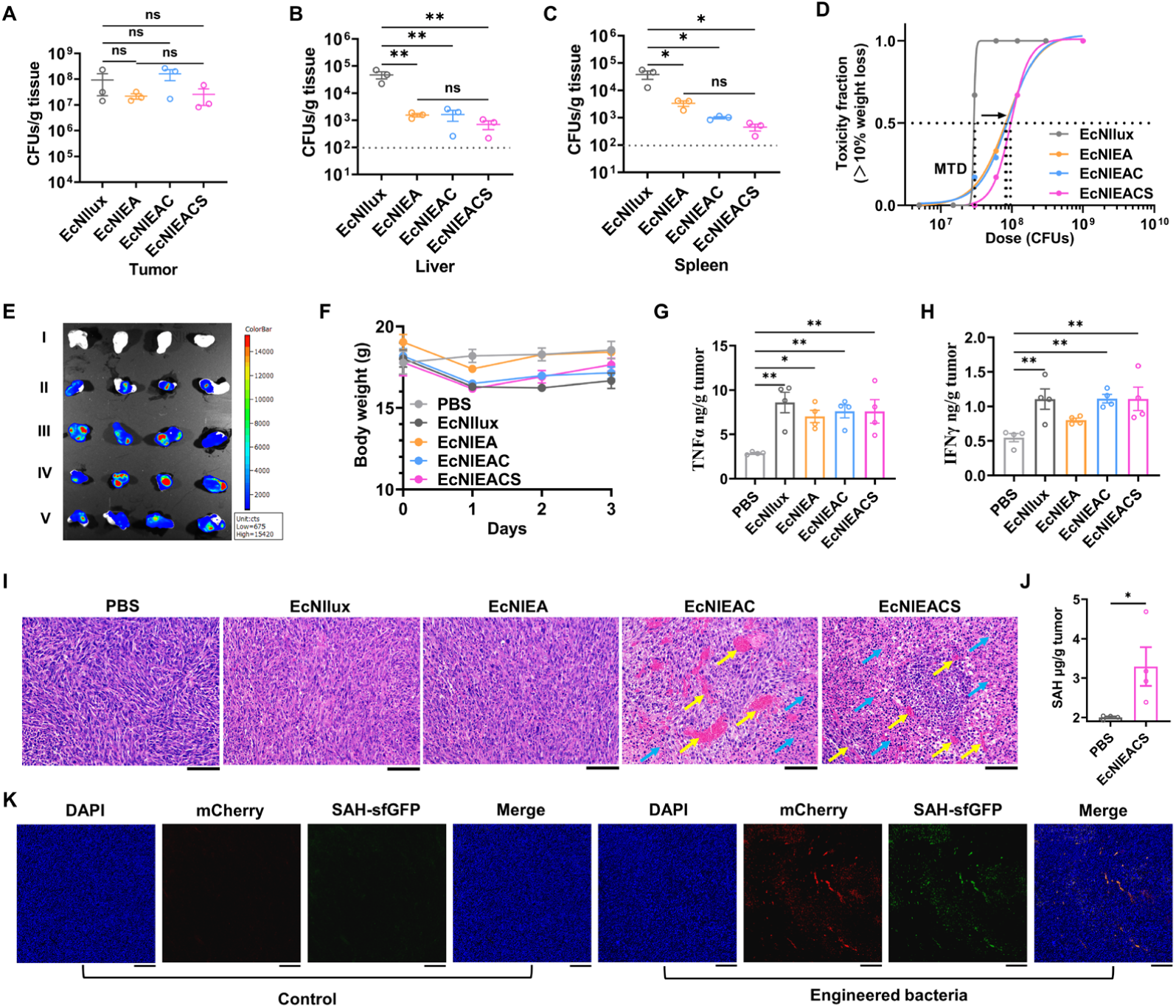
Characterization of engineered EcN functions in tumors. (A-C) Quantification of bacterial colonization in tumors and different organs harvested from mice bearing the MC38 tumor after inject bacterial for 24 h. LOD = 1 × 10^2^ CFU/g (mean ± SEM, n = 3). Statistical significance was determined by a one-way ANOVA, ns p > 0.05, *p ≤ 0.05, **p ≤ 0.01. (D) Dose-toxicity curve with MTD=2.9 × 10^7^ CFU, 8.5 × 10^7^ CFU, 8.9 × 10^7^ CFU and 9.7 × 10^7^ CFU for EcNllux, EcNlEA, EcNlEAC and EcNlEACS, respectively. MTD was calculated based on TD_50_ (n ≥ 6). (E) Visualization of bacterial colonization through bioluminescence imaging. I: PBS, II: EcNllux, III: EcNlEA, IV: EcNlEAC and V: EcNlEACS. (F) Weight changes in MC38-bearing mice during different strains treatment (mean ± SEM, n = 4). (G-H) Changes in intracellular cytokine (TNFα and IL-6) levels after 24 h of bacterial injection (mean ± SEM, n=4). Statistical significance was determined by a one-way ANOVA, *p≤ 0.05, **p ≤ 0.01. (I) Histological images of tumor sections from each group stained with hematoxylin and eosin. Yellow arrow: coagulation. Blue arrow: necrosis. Scale bars=100 μm. (J) Detection of intra-tumoral SAH levels after intravenous injection of EcNlEACS for 24 h (mean ± SEM, n = 4). Statistical significance was determined by an unpaired two-tailed t test, *p ≤ 0.05. (K) Fluorescence imaging of tumor slices. DAPI staining (blue), engineered bacteria (red), SAH (green). Scale bars = 200 μm.

EcNllux, EcNlEA, EcNlEAC and EcNlEACS, respectively.) in MC38 tumor-bearing mice, and the tumors were collected to measure the levels of cytokine by ELISA after 24 h injection. The results showed that all bacterial groups could significantly increase TNFα and IL-6 levels in tumors (Fig. 6G, H), which indicated that EcN could effectively stimulate the immune system. As assessed by H&E staining, treatment with EcNlEAC and EcNlEACS triggered apparent coagulation (yellow arrow), while this phenomenon was not observed in residual groups (Fig. 6I). Though EcNlEACS and EcNlEAC could cause tumor tissue damage, treatment with EcNlEACS exhibited a greater degree of necrosis (Fig. 6I). This result was due to the production of a large amount of SAH by EcNllEACS (Fig. 6J), and SAH did not leak outside the tumor tissues (Fig. S9).

To observe the expression of SAH in tumors visually, we fused the *sfGFP* tag with *SAH* gene. In addition, a *mCherry* gene was also integrated to observe the localization of EcNlEACS in tumors. Fluorescence scanning images showed that the engineered EcN effectively colonize tumors and express SAH after 24 h injection, but the PBS group did not display any red or green fluorescence signals (Fig. 6K). In summary, we demonstrated that EcNlEACS can perform the multiple functions in tumors: i) Specific and selective colonization in tumors. ii) Expression of coagulase to cause thrombus. iii) Self-adjusting release of cytotoxic proteins. These results provided further motivation for utilizing engineered EcN to treat tumors.

### Therapeutic Effects in Vivo

To assess the therapeutic potential of this system in vivo, we administrated MC38 tumor-bearing mice with PBS, EcNllux, EcNlEA, EcNlEAC and EcNlEACS for three times over the course of 6 days and monitored for 10 days (Fig. 7A). Although all bacterial administration groups resulted in a delay in tumor growth, EcNlEACS treatment showed the best anti-tumor among all groups. Administration of EcNlEACS could significantly inhibit tumor growth (Fig. 7B), and exhibited the lightest tumor weight (Fig. 7C), the smallest tumor size (Fig. 7D) and the lowest tumor cell proliferation activity (Fig. 7E). The tumor growth inhibition value was about 90% in 10^th^ day of EcNlEACS. Notably, all bacterial groups exhibited slight weight loss after 24 h of intravenous administration, and regained their weights at the end of 10 days (Fig. S10). Moreover, EcNlEACS treatment significantly prolonged the survival time of the MC38 tumor-bearing mice to a median survival time of 26 days compared to PBS (13 days), EcNllux (14 days), EcNlEA (15 days) and EcNlEAC (16 days) (Fig. 7F). All these results demonstrated the potential of our multifunctional live bacterial therapy to combat cancers.

**Fig. 7.**
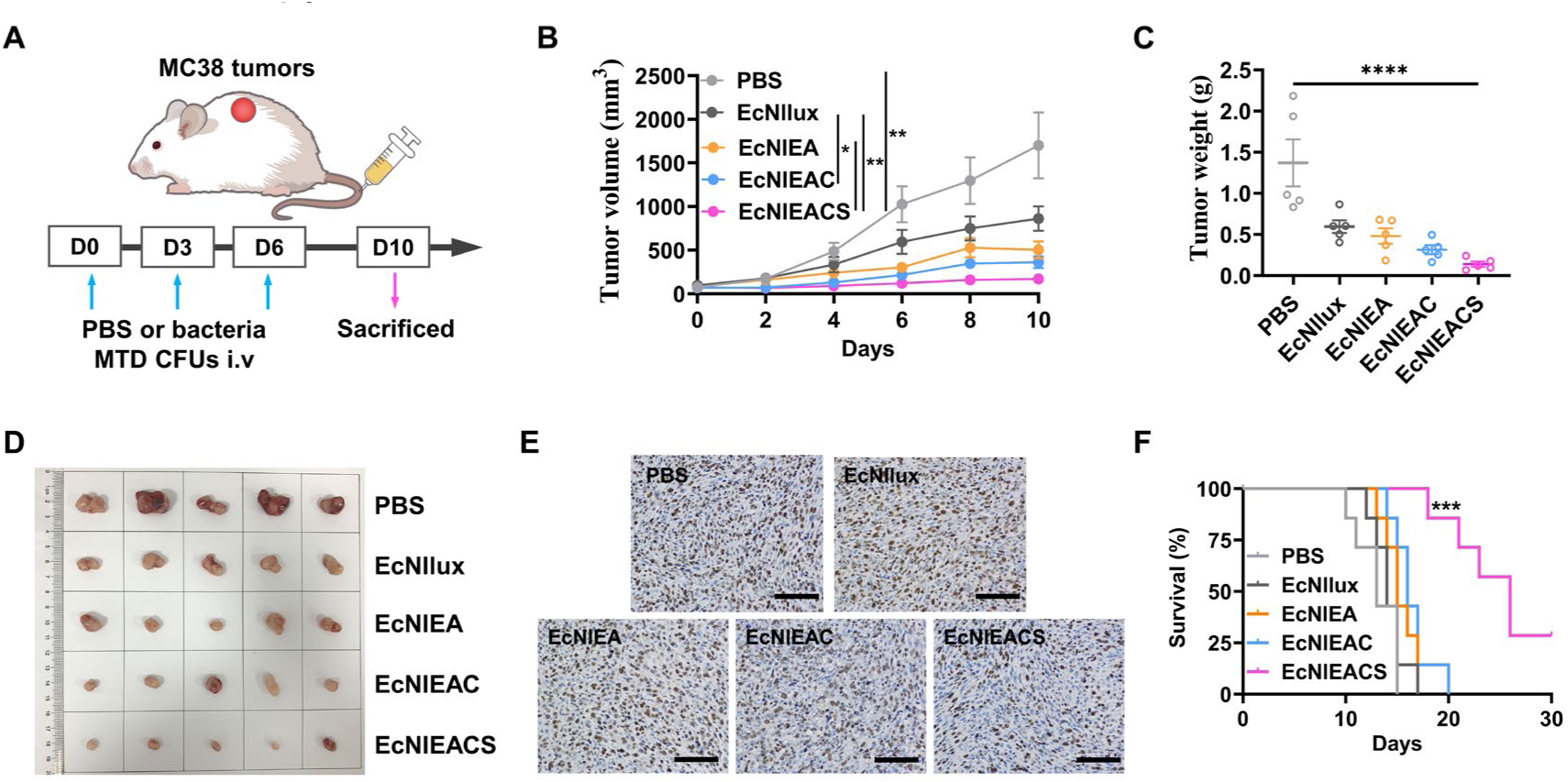
Evaluation of therapeutic efficacy of intravenous injection engineered EcN in a MC38 subcutaneous tumor model. (A) Therapeutic schedule. (B) Tumor growth of mice in different treatment groups within 10 days (mean ± SEM, n = 5). Statistical significance was determined by a two-way ANOVA, *p ≤ 0.05, **p ≤ 0.01. (C) Tumor weights of mice with different treatments on the 10^th^ day (mean ± SEM, n = 5). Statistical significance was determined by a one-way ANOVA, ****p ≤ 0.0001. (D) Images of the excised tumor tissues. (E) Immunohistochemistry analysis of MC38 tumor sections stained with Ki67 in PBS controls and groups treated with engineered EcN (mean ± SEM, n = 5). Scale bars= 100 μm. The sections were obtained from tumors showed in (D). (F) Survival curves for the mice receiving different treatments (mean ± SEM, n = 7). Statistical significance was determined by a Log-rank (Mantel-Cox) test, ***p ≤ 0.001.

## DISCUSSION

Here, we used synthetic biology strategies to design and create an intelligent genetic engineered probiotic strain EcNlEACS capable of selective tumor colonization, causing coagulation and self-adjusting drug delivery without additional inducers to fight tumors. The targeted colonization and self-seclusion in tumor sites were rationally designed by developing a lactate-sensing system to control the expression of growth essential gene *asd* and coagulase simultaneously. Subsequently, the QS system guaranteed to release therapeutic protein only in tumor and kill cancer cells.

As known, there is a significant difference in the content of lactate between normal tissues and tumor, which requires the sensing system to function at tumor concentrations strictly to avoid activation in other normal organs. One critical design of this gene circuit is optimizing a lactate sensing system based on LldR from *C. glutamicum* that is not inhibited by glucose. This system can respond to tumor only when the lactate concentration is above 5 mM, which is crucial for application in tumor treatment. We then coupled the optimized lactate biosensor with bacterial growth through the expression of an essential gene *asd* to address the limitation of targeted colonization in tumor. Moreover, the fusion of the *asd* gene with strong degradation label can further reduce the risk of bacterial escape. Induction of thrombosis in tumor vasculature is a promising avenue for the treatment of tumors. Unlike tumor embolism induced by pure thrombin, coagulase, or pathogenic bacteria, we used a tumor biomarker to regulate the expression of coagulase in a probiotic strain resulting in more targeted coagulation. The resultant enclosed environment not only helps to deprive tumor cells of oxygen and nutrients, which leads to tumor cell death, but also reduces the risk of EcNlEACS and therapeutic protein leakage. The other critical design of this gene circuit is exploiting quorum-sensing system for evoking the expression of therapeutic protein. Only when the engineered bacteria colonize the tumor and grow to a specific density, the expression of toxic proteins can be activated. Collectively, compared to other tumor treatment strategies that utilize bacteria as carriers, our multifunctional collaborative strategy enhanced treatment efficacy and improved biosafety.

A various bacterial based therapies have previously been investigated for the treatment of cancer, and attenuated pathogens such as *Salmonella typhimurium*, *Clostridium novyi*, and *Listeria monocytogenes* were widely used^5–7^. However, these pathogens have dose-limiting toxicities and are prone to causing infection once colonized in normal organs. In this study, we used a probiotic strain EcN as chassis for its advantageous safety and convenience of genetic manipulation. In addition, increasing the dosage without increasing side effects endows EcN with stronger therapeutic potential. Nonetheless, there are still some issues needing to be further resolved, such as how to reduce the immunogenicity of EcN during delivery to improve efficiency. Using biomaterial strategy is a reasonable option to improve the delivery of therapeutic bacteria, such as wrapping with cell membranes and programmable encapsulation system^35–37^. In short, we provided a strategy for effectively targeting and colonizing tumors which laying the foundation for enhancing maximum drug administration, and developed a treatment model with the “closed door” (tumor embolism) and the “released dogs” (therapeutic drugs) to achieve the goal of efficiently and safely killing cancer cells.

## ACKNOWLEDGMENTS

This work was sponsored by the National Natural Science Foundation of China (22134003), the National Key Research and Development Program of China (2020YFA0908800 and 2023YFF1204500) and China Postdoctoral Science Foundation (2023M741176).

## AUTHOR CONTRIBUTIONS

Z.-P.Z., Y.Z., and B.-C.Y. designed the experiments, wrote the paper and provided funding support. Z.-P.Z., J.M., and X.-P.Z. performed in vitro experiments. Z.-P.Z., X.-G.W., and S.-T.S. performed in vivo experiments. B.-C.Y. helped with manuscript revision.

## MATERIALS AND METHODS

### Ethics

Female C57BL/6 mice (5-6 weeks) were obtained from Shanghai Model Organisms Center, Inc. All animal procedures were performed in accordance with the Guidelines for Care and Use of Laboratory Animals of East China University of Science and Technology and the experiments were approved by the Animal Ethics Committee of East China University of Science and Technology. All animals were euthanized when tumor burden reaches 2000 mm^3^.

### Materials

Anti-α-hemolysin antibodies (ab190467) were purchased from Abcam. HRP conjugated anti-mouse IgG (HS201) and the cell counting kit-8 assay (CCK-8) reagent were purchased from TransGen Biotech. Mouse TNF-α ELISA kit (EMC102aQT), mouse IFN-γ ELISA kit (EMC101gQT) and mouse IL-6 ELISA kit (EMC004QT) were purchased from Neobioscience. HE dye solution set (G1003) was purchased from Servicebio. DNA polymerase (P505) was purchased from Vazyme. Blood agar plates (CP0230) were purchased from HuanKai Biology. Roswell Park Memorial Institute (RPMI) 1640 medium and Dulbecco’s modified Eagle’s medium (DMEM) were purchased from Gibco. DAP (BCCH0213) was purchased from Sigma-Aldrich. Human blood was obtained from Shanghai Tenth People’s Hospital.

### Bacteria and cell culture

All Bacterial strains and cell lines are listed in Table S1. Bacterial strains were spread on the LB solid plate (with appropriate antibiotics) and then incubated at 37°C for 12 h. Colonies were picked out and transferred to 700 μL LB liquid medium (with appropriate antibiotics) in a microplate shaker (37°C, 1000 rpm) for overnight. Afterward, the activated cultures were diluted 200-fold into fresh medium and further cultured to the logarithmic growth stage (OD≈0.4).

Cancer cells were maintained at 37°C in a 5% CO_2_ humidified atmosphere. MC38, SW480, B16 cells were cultured in cell culture flasks containing 5 mL of 1640 medium, respectively, supplemented with 10% fetal bovine serum (FBS) and 100 U/mL penicillin-streptomycin. MCF-7 cells were cultured in DMEM medium with 10% fetal bovine serum (FBS) and 100 U/mL penicillin-streptomycin.

### Construction of plasmids

The construction of all plasmids was performed in *E. coli* strain DH5α (TransGen Biotech) in accordance with standard procedures. Plasmids were constructed by an origin replication of pSC101. To construct and optimize lactate-responsive gene circuits, the codon-optimized *lldR* gene (Sangon Biotech) from *C. glutamicum* was placed downstream of a series of different combinations of promoters and RBSs and the expression of the *sfGFP* gene was regulated by a synthetic promoter, which containing two LldR repressor binding sites. To construct biocontainment circuits, the essential gene *asd* or its variants (asdaav and asdlaa) was added downstream of the synthesis promoter regulated by L-lactate. The *asd* gene was obtained by colony polymerase chain reaction (PCR) from *E. coli* Nissle 1917, and the protein-degradation tag of AAV or LAA was added to the C-terminus of the *asd* gene through PCR. Codon-optimized *SAH* and truncated *coaA* gene were integrated into engineered plasmid through homologous recombination. Standard strength promoter (BBa_J231XX) sequences were obtained from http://parts.igem.org. All sequences are shown in Table S2.

### *asd* gene deletion and *luxE* gene integration in EcNl strain

The deletion of the *asd* gene and the integration of the *luxE* gene were carried out through the λ-Red recombination system. Details of chromosomal gene deletion and integration were previously reported^38, 39^. In brief, linear DNA containing a P_j23108_-*luxE*, an aminoglycoside phosphotransferase-gene (*kanR*) gene and two homology arms (50 bp) was amplified by PCR, and electroporated into EcNl strain carrying pKD46 plasmid. Bacteria were spread on a LB solid plate with 100 μg/mL DAP and cultured for overnight. Chromosomal deletions and integration were verified by colony PCR and sequencing.

### Determination of fluorescence intensity

Each activated bacterial strain was diluted 200-fold into fresh LB liquid medium containing different concentrations of L-lactate in a microplate shaker (1000 rpm, 37°C). After 12 h, the fluorescence values and OD_600_ were measured. The Fluo/OD_600_ showed the reporter sfGFP expression intensity. It is noted that before fluorescence detection, the cultures were needed to be washed twice and resuspended in PBS. OD_600_ and fluorescence values were measured by a microplate reader (BioTek Instruments, Winooski, VT, USA).

### Bacterial escape analysis

Activated bacteria were washed twice by PBS and diluted 1000-fold into 200 μL fresh LB liquid medium containing different concentrations of L-lactate or 100 μg/mL DAP. After 8 h cultivation at 37°C, 1000 rpm, all cultures were serially diluted and spread on the LB solid plate with 200 μg/mL DAP, after which colonies were counted after 12 h culture. The escape rate was defined as the ratio between colonies grown in DAP and different concentrations of L-lactate.

### Expression and purification of truncated CoaA and SAH protein

The *coaA* or *SAH* gene containing the 6 × His tag was inserted into a pET-28a expression vector respectively. The protein was expressed and purified according to standard protocols in *E. coli* BL21 strain.

### Blood clotting in vitro

For pure enzymes assay, 0, 0.15 μg and 1.5 μg coagulase (CoaA) were incubated with 50 μL human blood at 37°C, respectively. Photos were taken at 30 min and 60 min. For engineered EcN assay, bacterial strains were cultured to OD_600_≈0.4, then induced by different concentrations of L-lactate for 6 h. Afterward, these bacteria were incubated with human blood at 37°C, respectively. Photos were taken at 3 h and 12 h.

### Western blot analysis for SAH

Activated EcNlEAC and EcNlEACS strains were diluted 200-fold into fresh LB liquid medium with 100 μg/mL streptomycin and DAP, and grown in a 37 °C shaker for 12h. Cultures were collected and centrifuged at 12000 rcf for 5 min at 4°C. The supernatant was used for Western blot (WB) analysis. Supernatant and LB control were loaded per well for SDS-PAGE, followed by transferring to nitrocellulose membranes and WB test was the performed according to a standard protocol, with an appropriate concentration of SAH antibody (1:2000) and a HRP conjugated goat anti-mouse IgG (1:5000).

### ELISA analysis for SAH

Activated EcNlEACS strain was diluted 200-fold into fresh LB liquid medium with 100 μg/mL streptomycin and DAP, and grown in a 37°C shaker for 12 h. Cultures were collected at 6 h, 8 h, 10 h, 12 h, and then centrifuged at 12000 rcf for 5 min at 4°C. The supernatant was used for ELISA analysis. The standard curve was obtained through purified SAH protein. The SAH protein was diluted to 100, 50, 10, 5, 1 and 0 ng/mL. 100 μL diluted SAH was added to per well of ELISA plate and incubated overnight at 4°C, then the well plate was washed three times with 250 μL TBST buffer. After blocking with 5% bovine serum albumin (200 μL) for 1 h at 37°C and subsequent washing step, 100 μL of SAH antibody (1:2000) was added to each well, incubated for 1 h at 37°C and subjected to subsequent washing step. HRP conjugated goat anti-mouse IgG (1:3000) was added to each well, incubated for 1 h at 37°C and subsequently washed. After the washing step, 200 μL of TMB was added to each well, and the plate was incubated for 15 min at 37°C. 2 M sulfuric acid was added to each well to stop the reaction, and the OD_450_ was then measured.

### Cytotoxicity in vitro

For CCK-8 assay, MCF-7 cells (1×10^4^ cells/100 μL) were seeded into a 96-well plate and for 12 h, then 10 μL PBS, 50 ng/mL SAH protein, 10 μL 10% 10× LB medium and 10 μL 10% 10× concentrated, sterile supernatant from EcNlEAC and EcNlEACS were added to culture medium for 6 h. After that, samples were washed two times with 100 μL of PBS, followed by the addition of 100 μL of RPMI-1640 medium with 10% CCK-8 reagent. After 20 min incubation at 37°C, the OD_450_ values of all samples were recorded. Cell viability (%) = (OD_Sample_ - OD_Blank_) / (OD_PBS_ - OD_Blank_) × 100. OD_Blank_: The OD_450_ of 1640 medium without CCK8 reagent.

### Quantification of bacterial colonization

Overnight cultures of all strains were grown in LB liquid medium with 100 μg/mL streptomycin and 100 μg/mL DAP. A 1/200 dilution was made by fresh LB with 100 μg/mL streptomycin and 100 μg/mL DAP on the following day and grown in a 37 °C shaker until OD_600_≈0.4. Each sample was washed two times with cold PBS, and then diluted to 1.5×10^8^ CFU/mL in PBS. Mice were injected intravenously with 100 μL diluted bacteria and sacrificed at a preset time (24 h or 48 h). Tumors and organs were immerged in 500 μL of PBS and homogenized using a tissue homogenizer. The homogenized samples were diluted and spread on the LB agar plate, containing 100 μg/mL DAP, after which colonies were counted after 12 h. The bacterial titer (CFU/g tissue) was calculated according to colony counts, dilution ratio and the tissue weight.

### H&E and immunohistochemistry

Sample sections were fixed on slides (5 μm) and stained with hematoxylin and eosin. The immunohistochemistry was completed by Servicebio. Photos were taken by an inverted microscope.

### Detection of serum transaminases and cytokines

Each bacterial strain or PBS was intravenously injected to female C57BL/6 mice. After 24 or 48 h, the blood samples were collected and centrifuged at 2000 rcf for 10 min at 4°C. The serum was collected and stored at −80°C for analysis. Biochemical indicators were detected through a biochemical analyzer. Cytokines were measured using a ELISA kit according to the manufacturer’s protocol.

### Intra-tumoral cytokine and SAH analysis

Each bacterial strain or PBS was intravenously injected to MC38-bearing female C57BL/6 mice. After 24 h, Tumors were collected and homogenized in 500 μL of PBS. The samples were centrifuged at 12000 rcf for 10 min at 4 °C. The supernatant was collected for the detection of cytokine and SAH by ELISA.

### Animal model and treatment assays

After a week of adaptive feeding, female C57BL/6 mice were subcutaneously injected with 5 × 10^5^ MC38 cells, 100 μL. For tumor treatment, tumors were grown to a volume was of 50-100 mm^3^ before bacteria or PBS injection, and bacteria and PBS were intravenously injected three times over the course of 6 days (on 0^th^, 3^th^, and 6^th^ day). Tumor volume (mm^3^) was measured with the caliper and calculated using the following formula: (length × width^2^)/2.

## REFERENCES

1. Petroni, G., Cantley, L.C., Santambrogio, L., Formenti, S.C. & Galluzzi, L. Radiotherapy as a tool to elicit clinically actionable signalling pathways in cancer. Nat. Rev. Clin. Oncol. 19, 114–131 (2022).

2. Galluzzi, L., Bravo-San Pedro, J.M., Demaria, S., Formenti, S.C. & Kroemer, G. Activating autophagy to potentiate immunogenic chemotherapy and radiation therapy. Nat. Rev. Clin. Oncol. 14, 247–258 (2017).

3. Waldman, A.D., Fritz, J.M. & Lenardo, M.J. A guide to cancer immunotherapy: from T cell basic science to clinical practice. Nat. Rev. Immunol. 20, 651–668 (2020).

4. Flugel, C.L. et al. Overcoming on-target, off-tumour toxicity of CAR T cell therapy for solid tumours. Nat. Rev. Clin. Oncol. 20, 49–62 (2023).

5. Zhou, S., Gravekamp, C., Bermudes, D. & Liu, K. Tumour-targeting bacteria engineered to fight cancer. Nat. Rev. Cancer 18, 727–743 (2018).

6. Sieow, B.F.-L., Wun, K.S., Yong, W.P., Hwang, I.Y. & Chang, M.W. Tweak to Treat: Reprograming Bacteria for Cancer Treatment. Trends Cancer 7, 447–464 (2021).

7. Gurbatri, C.R., Arpaia, N. & Danino, T. Engineering bacteria as interactive cancer therapies. Science 378, 858–864 (2022).

8. O’Donnell, M.A. Optimizing BCG therapy. Urol. Oncol.-Semin. Orig. Investig. 27, 325–328 (2009).

9. Lin, Q. et al. IFN-γ-dependent NK cell activation is essential to metastasis suppression by engineered Salmonella. Nat. Commun. 12, 2537 (2021).

10. Lynch, J.P., Goers, L. & Lesser, C.F. Emerging strategies for engineering Escherichia coli Nissle 1917-based therapeutics. Trends Pharmacol. Sci. 43, 772–786 (2022).

11. Jiang, S.-N. et al. Engineering of Bacteria for the Visualization of Targeted Delivery of a Cytolytic Anticancer Agent. Mol. Ther. 21, 1985–1995 (2013).

12. Ryan, R.M. et al. Bacterial delivery of a novel cytolysin to hypoxic areas of solid tumors. Gene Ther. 16, 329–339 (2009).

13. Wei, C. et al. Bifidobacteria Expressing Tumstatin Protein for Antitumor Therapy in Tumor-Bearing Mice. Technol. Cancer Res. Treat. 15, 498–508 (2015).

14. Chiang, C.-J. & Hong, Y.-H. In situ delivery of biobutyrate by probiotic Escherichia coli for cancer therapy. Sci. Rep. 11, 18172 (2021).

15. Gu, J. et al. Knockdown of HIF-1α by siRNA-expressing plasmid delivered by attenuated Salmonella enhances the antitumor effects of cisplatin on prostate cancer. Sci. Rep. 7, 7546 (2017).

16. Savage, T.M. et al. Chemokines expressed by engineered bacteria recruit and orchestrate antitumor immunity. Sci. Adv. 9, eadc9436 (2023).

17. St. Jean, A.T., Swofford, C.A., Panteli, J.T., Brentzel, Z.J. & Forbes, N.S. Bacterial Delivery of Staphylococcus aureus α-Hemolysin Causes Regression and Necrosis in Murine Tumors. Mol. Ther. 22, 1266–1274 (2014).

18. Tan, W. et al. Targeting of pancreatic cancer cells and stromal cells using engineered oncolytic Salmonella typhimurium. Mol. Ther. 30, 662–671 (2022).

19. Chen, Y., Du, M., Yuan, Z., Chen, Z. & Yan, F. Spatiotemporal control of engineered bacteria to express interferon-γ by focused ultrasound for tumor immunotherapy. Nat. Commun. 13, 4468 (2022).

20. Gurbatri, C.R. et al. Engineered probiotics for local tumor delivery of checkpoint blockade nanobodies. Sci. Transl. Med. 12, eaax0876 (2020).

21. Abedi, M.H. et al. Ultrasound-controllable engineered bacteria for cancer immunotherapy. Nat. Commun. 13, 1585 (2022).

22. Yu, B. et al. Explicit hypoxia targeting with tumor suppression by creating an “obligate” anaerobic Salmonella Typhimurium strain. Sci. Rep. 2, 436 (2012).

23. Chien, T. et al. Enhancing the tropism of bacteria via genetically programmed biosensors. *Nat*. Biomed. Eng 6, 94–104 (2022).

24. Zúñiga, A. et al. Engineered l-Lactate Responding Promoter System Operating in Glucose-Rich and Anoxic Environments. ACS Synth. Biol. 10, 3527–3536 (2021).

25. Seidi, K. et al. NGR (Asn-Gly-Arg)-targeted delivery of coagulase to tumor vasculature arrests cancer cell growth. Oncogene 37, 3967–3980 (2018).

26. de la Cruz-López, K.G., Castro-Muñoz, L.J., Reyes-Hernández, D.O., García-Carrancá, A. & Manzo-Merino, J. Lactate in the Regulation of Tumor Microenvironment and Therapeutic Approaches. Front. Oncol. 9 (2019).

27. Ho, J.M.L., Miller, C.A., Parks, S.E., Mattia, J.R. & Bennett, Matthew R. A suppressor tRNA-mediated feedforward loop eliminates leaky gene expression in bacteria. Nucleic Acids Res. 49, e25–e25 (2021).

28. Gao, Y.-G. et al. Structural and functional characterization of the LldR from Corynebacterium glutamicum: a transcriptional repressor involved in l-lactate and sugar utilization. Nucleic Acids Res. 36, 7110–7123 (2008).

29. Georgi, T., Engels, V. & Wendisch Volker, F. Regulation of l-Lactate Utilization by the FadR-Type Regulator LldR of Corynebacterium glutamicum. J. Bacteriol. 190, 963–971 (2008).

30. Zou, Z.-P., Yang, Y., Wang, J., Zhou, Y. & Ye, B.-C. Coupling split-lux cassette with a toggle switch in bacteria for ultrasensitive blood markers detection in feces and urine. Biosens. Bioelectron. 214, 114520 (2022).

31. Wang, Y. et al. Active recruitment of anti–PD-1–conjugated platelets through tumor-selective thrombosis for enhanced anticancer immunotherapy. Sci. Adv. 9, eadf6854 (2023).

32. Li, S. et al. A DNA nanorobot functions as a cancer therapeutic in response to a molecular trigger in vivo. Nat. Biotechnol. 36, 258–264 (2018).

33. Panizzi, P. et al. Fibrinogen Substrate Recognition by Staphylocoagulase·(Pro)thrombin Complexes*. J. Biol. Chem. 281, 1179–1187 (2006).

34. Dang, Z. et al. Synthetic bacterial therapies for intestinal diseases based on quorum-sensing circuits. Biotechnol. Adv. 65, 108142 (2023).

35. Cao, Z., Cheng, S., Wang, X., Pang, Y. & Liu, J. Camouflaging bacteria by wrapping with cell membranes. Nat. Commun. 10, 3452 (2019).

36. Harimoto, T. et al. A programmable encapsulation system improves delivery of therapeutic bacteria in mice. Nat. Biotechnol. 40, 1259–1269 (2022).

37. Wang, X. et al. Bioinspired oral delivery of gut microbiota by self-coating with biofilms. Sci. Adv. 6, eabb1952 (2020).

38. Zou, Z.-P., Du, Y., Fang, T.-T., Zhou, Y. & Ye, B.-C. Biomarker-responsive engineered probiotic diagnoses, records, and ameliorates inflammatory bowel disease in mice. Cell Host Microbe 31, 199–212.e195 (2023).

39. Zou, Z.-P. et al. Protocol for engineering E. coli Nissle 1917 to diagnose, record, and ameliorate inflammatory bowel disease in mice. STAR Protoc. 4, 102254 (2023).

