## Supplemental Figure 1 for "Self-adjusting Engineered Probiotic for Targeted Tumor Colonization and Local Therapeutics Delivery"

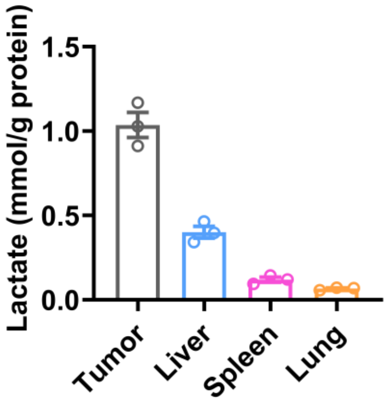


**Fig. S1. Lactate levels in MC38 tumors and healthy organs.** MC38 cells were injected at a volume of 100 μL ($5\times10^5$ cells). Tumours were grown to an average of approximately 150 $mm^3$ and then collected tumours and organs. Lactate concentrations were detected by lactate colorimetric assay kits (mean ± SEM, n=3).
