## Supplemental Figure 2 for "Self-adjusting Engineered Probiotic for Targeted Tumor Colonization and Local Therapeutics Delivery"

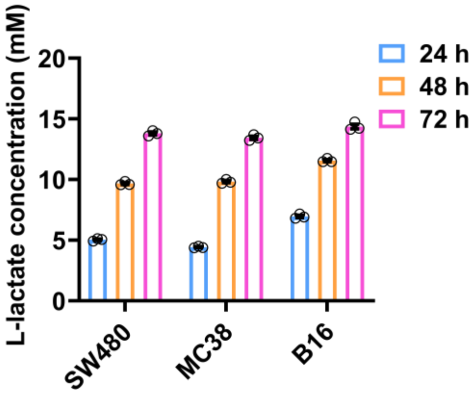


**Fig. S2. Lactate levels in various cancer cell lines.** Cell culture medium supernatant from four cancer cell lines (SW480, MC38, B16) was collected in 24 h, 48 h, 72 h. Lactate concentrations were detected by lactate colorimetric assay kits (mean ± SEM, n=3).
