## Supplemental Figure 3 for "Self-adjusting Engineered Probiotic for Targeted Tumor Colonization and Local Therapeutics Delivery"

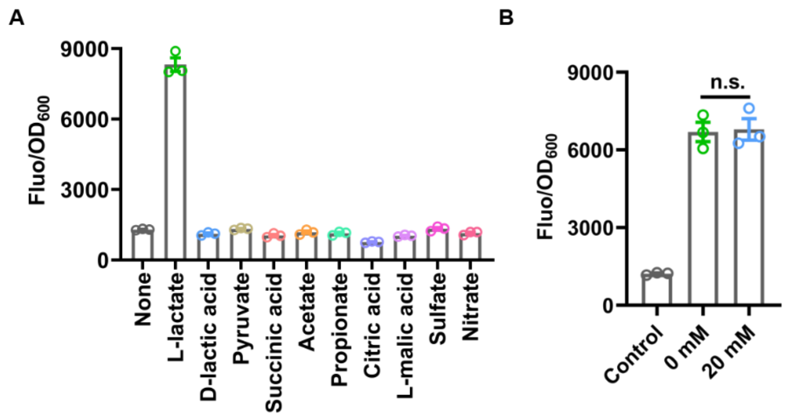


**Fig. S3. Specificity test and glucose inhibition effect analysis of the L-lactate responsive biosensor, L1032.** (A) The specificities of L1032 were determined, and all ligands were tested at a concentration of 10 mM (mean ± SEM, n=3). (B) L1032 was cultured in presence of 10 mM L-lactate and 10 or 20 mM glucose. No decrease in response indicates no inhibition by the added glucose (mean ± SEM, n=3).
