## Supplemental Figure 4 for "Self-adjusting Engineered Probiotic for Targeted Tumor Colonization and Local Therapeutics Delivery"

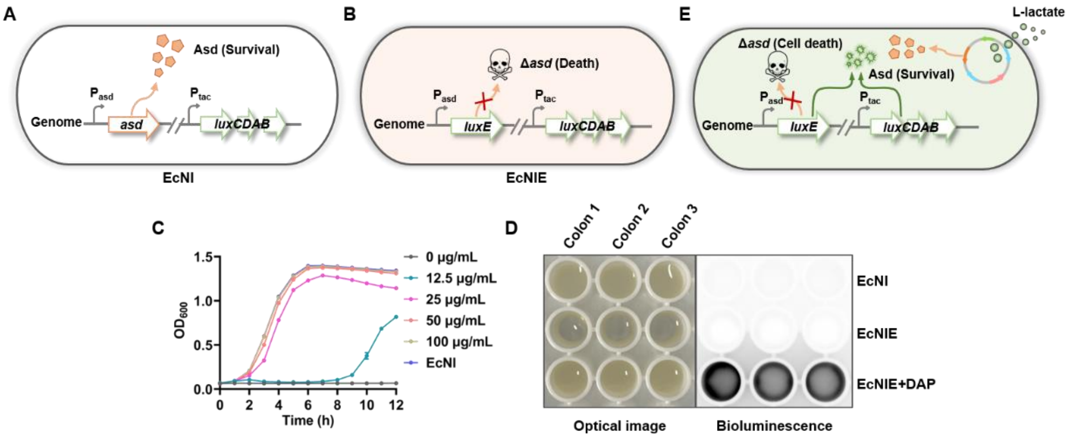


**Fig. S4. Construction and characterization of a EcN strain with the *asd* gene knockout.** (A) Schematic diagram of EcNI strain. (B) Schematic diagram of EcNIE strain, which the *asd* gene has been replaced by *luxE*. (C) Growth ability of EcNIE strain (mean ± SEM, n=3). (D) Visualization of EcNIE strain growth through optical image and bioluminescence. (E) Schematic diagram of a engineered EcN strain, which growth is regulated by L-lactate.
