## Supplemental Figure 5 for "Self-adjusting Engineered Probiotic for Targeted Tumor Colonization and Local Therapeutics Delivery"

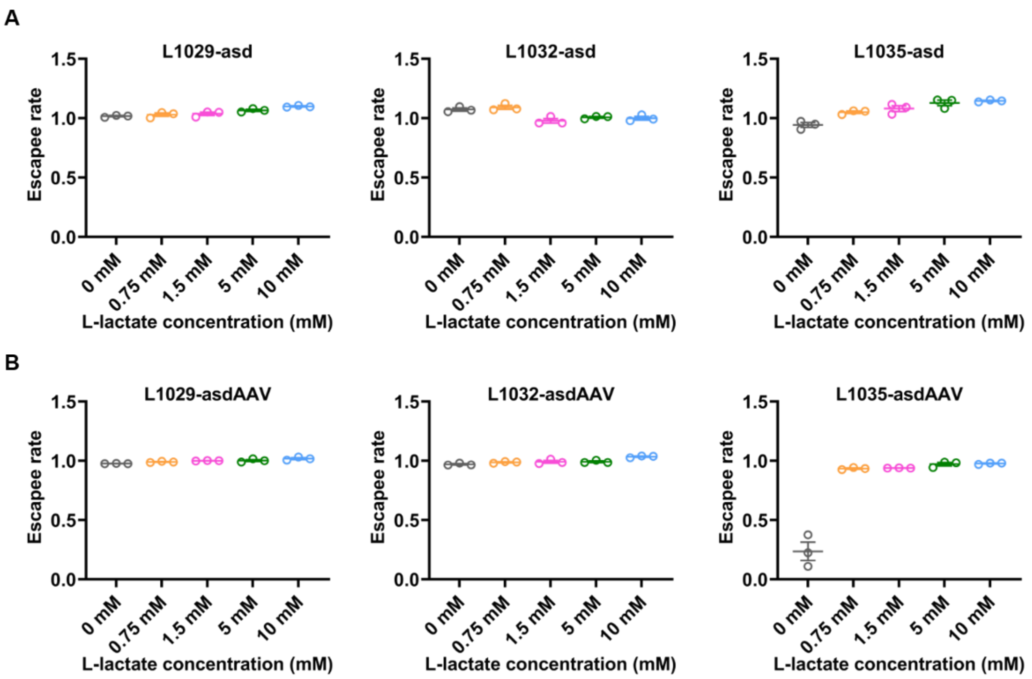


**Fig. S5. Coupling a L-lactate biosensor with bacterial growth via the expression of an essential gene *asd*.** Characterization of some biocontainment variants on the escapee rate (mean ± SEM, n=3).
