## Supplemental Figure 6 for "Self-adjusting Engineered Probiotic for Targeted Tumor Colonization and Local Therapeutics Delivery"

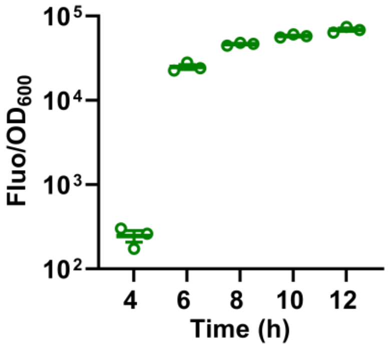


**Fig. S6. Characterization of quorum sensing elements.** The fluorescence signal of EcNl strain containing a quorum-sensing sensor increases with the increasing bacterial concentration (mean ± SEM, n=3).
