## Supplemental Figure 7 for "Self-adjusting Engineered Probiotic for Targeted Tumor Colonization and Local Therapeutics Delivery"

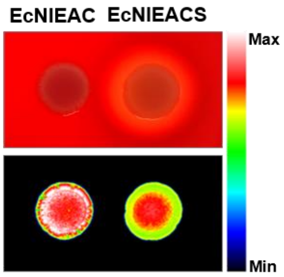


**Fig. S7. Visualization of EcNIEACS strain lysing red blood cells.** Add 10 μL EcNIAC ($10^9$ CFU) and EcNIACS ($10^9$ CFU) cultures to the surface of the blood agar plate, continue to culture at 37℃ for 12 h, and then image. Clear zone around the EcNIACS colonies indicating the expression of SAH.
