## Supplemental Figure 8 for "Self-adjusting Engineered Probiotic for Targeted Tumor Colonization and Local Therapeutics Delivery"

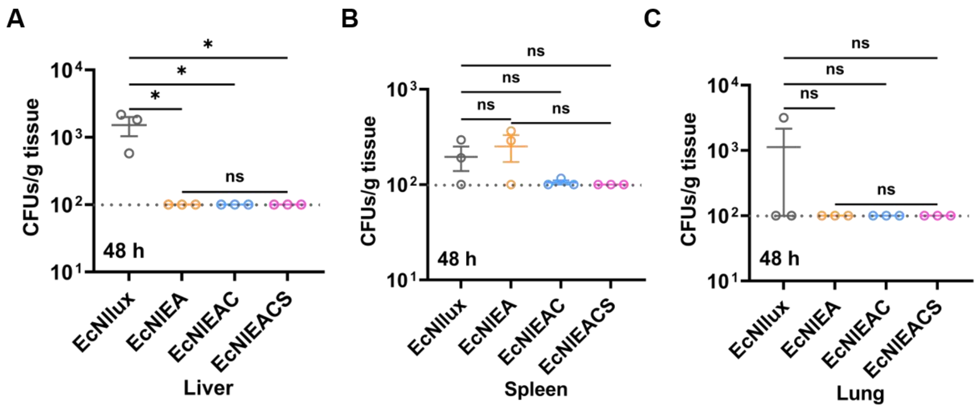


**Fig. S8. Selective colonization of engineered EcN.** Quantification of bacterial colonization in different organs (liver, spleen, lung) harvested from healthy mice after inject bacterial for 48 h. LOD=1×$10^2$ CFU/g (mean ± SEM, n=3).
