## Supplemental Figure 9 for "Self-adjusting Engineered Probiotic for Targeted Tumor Colonization and Local Therapeutics Delivery"

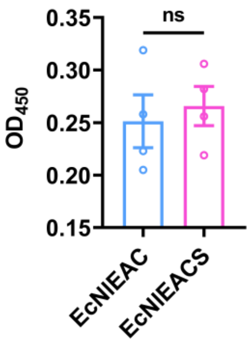


**Fig. S9. Detection of serum SAH levels after intravenous injection of EcNIEAC and EcNIEACS for 24 h.** Statistical significance was determined by an unpaired two-tailed t test, ns $p > 0.05$ (mean ± SEM, n = 4).
