## Supplemental Figure 10 for "Self-adjusting Engineered Probiotic for Targeted Tumor Colonization and Local Therapeutics Delivery"

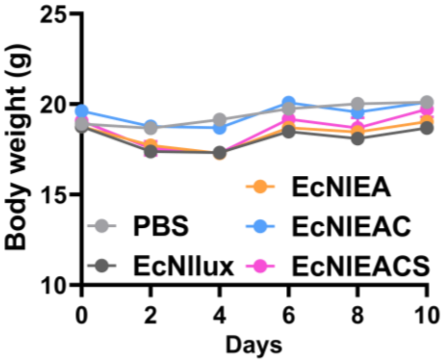


**Figure S10. Mouse weight change during treatment.** During the treatment process, the weight of the mice was measured every two days.
