## Supplemental Table 1 for "Self-adjusting Engineered Probiotic for Targeted Tumor Colonization and Local Therapeutics Delivery"

**Table S1** Summary of the strains used in this study

| Lable | Characteristics | promoter and RBS of *lldR* | Promoter and RBS of *sfGFP* | promoter of *luxAB* | promoter of *luxCD* | promoter of *luxE* | promoter and RBS of *asd* | promoter and RBS of *coaA* | promoter and RBS of *luxR* | RBS of *luxI* | Promoter of *SAH* |
|---|---|---|---|---|---|---|---|---|---|---|---|
| AL | *araC, lldR, sfGFP* | $P_{ara}$- $RBS_{ara}$ | $P_{tacO}$-B0035 | N/A | N/A | N/A | N/A | N/A | N/A | N/A | N/A |
| ALO | *araC, lldR, sfGFP* | $P_{ara}$- $RBS_{ara}$ | $P_{Otac}$-B0035 | N/A | N/A | N/A | N/A | N/A | N/A | N/A | N/A |
| ALOO | *araC, lldR, sfGFP* | $P_{ara}$- $RBS_{ara}$ | $P_{OtacO}$-B0035 | N/A | N/A | N/A | N/A | N/A | N/A | N/A | N/A |
| L0435 | *lldR, sfGFP* | $P_{j23104}$- B0035 | $P_{OtacO}$-B0035 | N/A | N/A | N/A | N/A | N/A | N/A | N/A | N/A |
| L0835 | *lldR, sfGFP* | $P_{j23108}$-B0035 | $P_{OtacO}$-B0035 | N/A | N/A | N/A | N/A | N/A | N/A | N/A | N/A |
| L1035 | *lldR, sfGFP* | $P_{j23110}$-B0035 | $P_{OtacO}$-B0035 | N/A | N/A | N/A | N/A | N/A | N/A | N/A | N/A |
| L1029 | *lldR, sfGFP* | $P_{j23110}$-B0029 | $P_{OtacO}$-B0035 | N/A | N/A | N/A | N/A | N/A | N/A | N/A | N/A |
| L1032 | *lldR, sfGFP* | $P_{j23110}$-B0032 | $P_{OtacO}$-B0035 | N/A | N/A | N/A | N/A | N/A | N/A | N/A | N/A |
| L0432 | *lldR, sfGFP* | $P_{j23104}$-B0032 | $P_{OtacO}$-B0035 | N/A | N/A | N/A | N/A | N/A | N/A | N/A | N/A |
| EcNI | EcN::*luxABCD* | N/A | N/A | $P_{tac}$ | $P_{tac}$ | $P_{j23108}$ | N/A | N/A | N/A | N/A | N/A |
| EcNIE | EcN::*luxABCDE Δasd* | N/A | N/A | $P_{tac}$ | $P_{tac}$ | $P_{j23108}$ | N/A | N/A | N/A | N/A | N/A |
| L1029asd | Use EcNIE as the chassis, *lldR, asd* | $P_{j23110}$-B0029 | N/A | $P_{tac}$ | $P_{tac}$ | $P_{j23108}$ | $P_{OtacO}$-B0035 | N/A | N/A | N/A | N/A |
| L1032asd | Use EcNIE as the chassis, *lldR, asd* | $P_{j23110}$-B0032 | N/A | $P_{tac}$ | $P_{tac}$ | $P_{j23108}$ | $P_{OtacO}$-B0035 | N/A | N/A | N/A | N/A |
| L1035asd | Use EcNIE as | $P_{j23110}$-B0035 | N/A | $P_{tac}$ | $P_{tac}$ | $P_{j23108}$ | $P_{OtacO}$- | N/A | N/A | N/A | N/A |

| | | | | | | | | | | | |
|---|---|---|---|---|---|---|---|---|---|---|---|
| | the chassis, *lldR, asd* | | | | | | B0035 | | | | |
| L1029asdAAV | Use EcNIE as the chassis, *lldR, asd-laa* | $P_{j23110}$-B0029 | N/A | $P_{tac}$ | $P_{tac}$ | $P_{j23108}$ | $P_{OtacO}$-B0035 | N/A | N/A | N/A | N/A |
| L1032asdAAV | Use EcNIE as the chassis, *lldR, asd-laa* | $P_{j23110}$-B0032 | N/A | $P_{tac}$ | $P_{tac}$ | $P_{j23108}$ | $P_{OtacO}$-B0035 | N/A | N/A | N/A | N/A |
| L1035asdAAV | Use EcNIE as the chassis, *lldR, asd-laa* | $P_{j23110}$-B0035 | N/A | $P_{tac}$ | $P_{tac}$ | $P_{j23108}$ | $P_{OtacO}$-B0035 | N/A | N/A | N/A | N/A |
| L1029asdLAA | Use EcNIE as the chassis, *lldR, asd-laa* | $P_{j23110}$-B0029 | N/A | $P_{tac}$ | $P_{tac}$ | $P_{j23108}$ | $P_{OtacO}$-B0035 | N/A | N/A | N/A | N/A |
| L1032asdLAA (EcNIEA) | Use EcNIE as the chassis, *lldR, asd-laa* | $P_{j23110}$-B0032 | N/A | $P_{tac}$ | $P_{tac}$ | $P_{j23108}$ | $P_{OtacO}$-B0035 | N/A | N/A | N/A | N/A |
| L1035asdLAA | Use EcNIE as the chassis, *lldR, asd-laa* | $P_{j23110}$-B0035 | N/A | $P_{tac}$ | $P_{tac}$ | $P_{j23108}$ | $P_{OtacO}$-B0035 | N/A | N/A | N/A | N/A |
| L1035coaA | Use EcNIE as the chassis, *lldR, coaA* | $P_{j23110}$-B0035 | N/A | $P_{tac}$ | $P_{tac}$ | $P_{j23108}$ | N/A | $P_{OtacO}$-B0035 | N/A | N/A | N/A |
| L1032asdcoaA | Use EcNIE as the chassis, *lldR, asd-laa*, | $P_{j23110}$-B0032 | N/A | $P_{tac}$ | $P_{tac}$ | $P_{j23108}$ | $P_{OtacO}$-B0035 | $P_{OtacO}$-B0035 | N/A | N/A | N/A |

| | | | | | | | | | | | |
|---|---|---|---|---|---|---|---|---|---|---|---|
| | *coaA* | | | | | | | | | | |
| EcNIEAC | Use EcNIE as the chassis, *lldR, asd-laa*, *coaA* | $P_{j23106}$-B0035 | N/A | $P_{tac}$ | $P_{tac}$ | $P_{j23108}$ | $P_{OtacO}$-RBS36 | $P_{OtacO}$-B0029 | N/A | N/A | N/A |
| EcNIEACS | Use EcNIE as the chassis, *lldR, asd-laa*, *coaA, luxI, luxR, SAH* | $P_{j23106}$-B0035 | N/A | $P_{tac}$ | $P_{tac}$ | $P_{j23108}$ | $P_{OtacO}$-RBS36 | $P_{OtacO}$-B0029 | $P_{j23108}$-B0035 | $RBS_{luxI}$ | $P_{lux}$ |
