## Supplemental Table 2 for "Self-adjusting Engineered Probiotic for Targeted Tumor Colonization and Local Therapeutics Delivery"

**Table S2** Sequences

| Name | Sequences |
|---|---|
| $P_{ara}$ | aaagccatgacaaaaacgcgtaacaaaagtgtctataatcacggcagaaaagtccacattgattatttgcacggcgtcacactttgctatgccatagcatttttatccataagattagcggatcctacctgacgctttttatcgcaactctctactgtttctccat |
| $P_{j23104}$ | ttgacagctagctcagtcctaggtattgtgctagc |
| $P_{j23108}$ | ctgacagctagctcagtcctaggtataatgctagc |
| $P_{j23110}$ | tttacggctagctcagtcctaggtacaatgctagc |
| $P_{j23106}$ | tttacggctagctcagtcctaggtatagtgctagc |
| $P_{tac}$ | ttgacaattaatcatcggctcgtataatg |
| $P_{tacO}$ | ttgacaattaatcatcggctcgtataatgcttgtggtctgaccaatga |
| $P_{Otac}$ | ttgacaatttggtctgaccacgtataatg |
| $P_{OtacO}$ | ttgacaatttggtctgaccacgtataatgcttgtggtctgaccaatga |
| $P_{lux}$ | agcacctgtaggatcgtacaggtttacgcaagaaaatggtttgttatagtcgaatgaattcattaaagaggagaaaggtacc |
| $RBS_{ara}$ | acccgtttttttgggaattcgagctctaaggaggttataaaaa |
| B0029 | tctagagttcacacaggaaacctactag |
| B0032 | tctagagtcacacaggaaagtactag |
| B0035 | tctagagattaaagaggagaaatactag |
| RBS36 | ctagtatttctcctgtgtgaactctaga |
| $RBS_{luxI}$ | gaattcattaaagaggagaaaggtacc |
| lldRO | tggtctgacca |
| *aav* | gctgcaaacgacgaaaactacgctgccgcagtt |
| *laa* | gctgcaaacgacgaaaactacgctttagccgca |
| *lldR* | atgagcgttaaagcgcatgaaagcgtgatggattgggttaccgaagaactgcgcagcggtcgcctgaaaattggtgatcatctgccgagcgaacgtgccctgagcgaaacactgggtgtgagccgtagcagcctgcgtgaagcactgcgtgtgctggaagcactgggtactattagcaccgccacgggcagcggtccacgtagtggtacaattattaccgcagcaccgggtcaggcactgagcctgagtgttaccctgcagctggttacgaatcaggttggtcatcatgatatttatgaaacccgtcagctgctggaaggttgggcagcactgcatagcagcgccgaacgtggtgattgggatgttgcggaagcactgctggaaaaaatggatgatccgaccctgccgctggaagattttctgcgttttgatgcggaatttcatgttgtcattagcaaaggtgcagaaaatccgctgattagtacactgatggaagcactgcgcctgagcgttgcagatcataccgttgcacgtgccctggccctgcctgattggcctgcaacaagcgcacgcctgc |

agaaagaacatcgtgccattctggccgctctgcgtgcaggtgaaagcacaatggcagcaaccctgattaaagatcatatcgaaggttattatcaggaaaccgcagcagcagaagcataa

*sfGFP* atgcgtaaaggcgaagagctgttcactggtgtcgtccctattctggtggaactggatggtgatgtcaacggtcataagttttccgtgcgtggcgagggtgaaggtgacgcaactaatggtaaactgacgctgaagttcatctgtacta
ctggtaaactgccggtaccttggccgactctggtaacgacgctgacttatggtgttcagtgctttgctcgttatccggaccatatgaagcagcatgacttcttcaagtccgccatgccggaaggctatgtgcaggaacgcacgatttc
ctttaaggatgacggcacgtacaaaacgcgtgcggaagtgaaatttgaaggcgataccctggtaaaccgcattgagctgaaaggcattgactttaaagaagacggcaatatcctgggccataagctggaatacaattttaaca
gccacaatgtttacatcaccgccgataaacaaaaaaatggcattaaagcgaattttaaaattcgccacaacgtggaggatggcagcgtgcagctggctgatcactaccagcaaaacactccaatcggtgatggtcctgttctgc
tgccagacaatcactatctgagcacgcaaagcgttctgtctaaagatccgaacgagaaacgcgatcatatggttctgctggagttcgtaaccgcagcgggcatcacgcatggtatggatgaactgtacaaatga

*luxA* atgaaatttggaaactttttgcttacataccaacctccccaattttctcaaacagaggtaatgaaacgtttggttaaattaggtcgcatctctgaggagtgtggttttgataccgtatggttactggagcatcatttcacggagtttggtttgct
tggtaacccttatgtcgctgctgcatatttacttggcgcgactaaaaaattgaatgtaggaactgccgctattgttcttcccacagcccatccagtacgccaacttgaagatgtgaatttattggatcaaatgtcaaaaggacgatttcg
gtttggtatttgccgagggctttacaacaaggactttcgcgtattcggcacagatatgaataacagtcgcgccttagcggaatgctggtacgggctgataaagaatggcatgacagagggatatatggaagctgataatgaacat
atcaagttccataaggtaaaagtaaaccccgcggcgtatagcagaggtggcgcaccggtttatgtggtggctgaatcagcttcgacgactgagtgggctgctcaatttggcctaccgatgatattaagttggattataaatactaac
gaaaagaaagcacaacttgagctttataatgaagtggctcaagaatatgggcacgatattcataatatcgaccattgcttatcatatataacatctgtagatcatgactcaattaaagcgaaagagatttgccggaaatttctgggg
cattggtatgattcttatgtgaatgctacgactatttttgatgattcagaccaaacaagaggttatgatttcaataaagggcagtggcgtgactttgtattaaaaggacataaagatactaatcgccgtattgattacagttacgaaatca
atcccgtgggaacgccgcaggaatgtattgacataattcaaaaagacattgatgctacaggaatatcaaatatttgttgtggatttgaagctaatggaacagtagacgaaattattgcttccatgaagctcttccagtctgatgtcatg
ccatttcttaaagaaaaacaacgttcgctattatattag

*luxB* atgaaatttggattgttcttccttaacttcatcaattcaacaactgttcaagaacaaagtatagttcgcatgcaggaaataacggagtatgttgataagttgaattttgaacagattttagtgtatgaaaatcatttttcagataatggtgttgt
cggcgctcctctgactgtttctggttttctgctcggtttaacagagaaaattaaaattggttcattaaatcacatcattacaactcatcatcctgtccgcatagcggaggaagcttgcttattggatcagttaagtgaagggagatttattta
gggtttagtgattgcgaaaaaaaagatgaaatgcattttttaatcgcccggttgaatatcaacagcaactatttgaagagtgttatgaaatcattaacgatgctttaacaacaggctattgtaatccagataacgatttttatagcttccc
taaaatatctgtaaatccccatgcttatacgccaggcggacctcggaaatatgtaacagcaaccagtcatcatattgttgagtgggcggccaaaaaaggtattcctctcatctttaagtgggatgattctaatgatgttagatatgaat
atgctgaaagatataaagccgttgcggataaatatgacgttgacctatcagagatagaccatcagttaatgatattagttaactataacgaagatagtaataaagctaaacaagagacgcgtgcatttattagtgattatgttcttga
aatgcaccctaatgaaaatttcgaaaataaacttgaagaaataattgcagaaaacgctgtcggaaattatacggagtgtataactgcggctaagttggcaattgaaaagtgtggtgcgaaaagtgtattgctgtcctttgaaccaa
tgaatgatttgatgagccaaaaaaatgtaatcaatattgttgatgataatattaagaagtaccacatggaatatacctaa

*luxC* atgactaaaaaaatttcattcattattaacggccaggttgaaatctttcccgaaggtgatgatttagtgcaatccattaattttggtgataatagtgtttacctgccaatattgaatgactctcatgtaaaaaacattattgattgtaatggaa
ataacgaattacggttgcataacattgtcaattttctctatacggtagggcaaagatggaaaaatgaagaatactcaagacgcaggacatacattcgtgacttaaaaaaatatatgggatattcagaagaaatggctaagctaga
ggccaattggatatctatgattttatgttctaaaggcggcctttatgatgttgtagaaaatgaacttggttctcgccatatcatggatgaatggctacctcaggatgaaagttatgttcgggcttttccgaaaggtaaatctgtacatctgttg
gcaggtaatgttccattatctgggatcatgtctatattacgcgcaattttaactaagaatcagtgtattataaaaacatcgtcaaccgatccttttaccgctaatgcattagcgttaagttttattgatgtagaccctaatcatccgataacg
cgctctttatctgttatatattggccccaccaaggtgatacatcactcgcaaaagaaattatgcaacatgcggatgttattgtcgcttggggagggccagatgcgattaattgggcggtagagcatgcgccatcttatgctgatgtgatt

aaatttggttctaaaaagagtctttgcattatcgataatcctgttgatttgacgtccgcagcgacaggtgcggctcatgatgtttgtttttacgatcagcgagcttgtttttctgcccaaaacatatattacatgggaaatcattatgaggaat

ttaagttagcgttgatagaaaaacttaatctatatgcgcatatattaccgaatgccaaaaaagattttgatgaaaaggcggcctattctttagttcaaaaagaaagcttgtttgctggattaaaagtagaggtggatattcatcaacgtt

ggatgattattgagtcaaatgcaggtgtggaatttaatcaaccacttggcagatgtgtgtaccttcatcacgtcgataatattgagcaaatattgccttatgttcaaaaaaaataagacgcaaaccatatctatttttccttgggagtcatc

atttaaatatcgagatgcgttagcattaaaaggtgcggaaaggattgtagaagcaggaatgaataacatatttcgagttggtggatctcatgacggaatgcgaccgttgcaacgattagtgacatatatttctcatgaaaggccatc

taactatacggctaaggatgttgcggttgaaatagaacagactcgattcctggaagaagataagttccttgtatttgtcccataa

*luxD* atggaaaatgaatcaaaatataaaaccatcgaccacgttatttgtgttgaaggaaataaaaaaattcatgtttgggaaacgctgccagaagaaaacagcccaaagagaaagaatgccattattattgcgtctggttttgcccgca

ggatggatcattttgctggtctggcggaatatttatcgcggaatggatttcatgtgatccgctatgattcgcttcaccacgttggattgagttcagggacaattgatgaatttacaatgtctataggaaagcagagcttgttagcagtggtt

gattggttaactacacgaaaaataaataacttcggtatgttggcttcaagcttatctgcgcggatagcttatgcaagcctatctgaaatcaatgcttcgtttttaatcaccgcagtcggtgttgttaacttaagatattctcttgaaagagctt

tagggtttgattatctcagtctacccattaatgaattgccgaataatctagattttgaaggccataaattgggtgctgaagtctttgcgagagattgtcttgattttggttgggaagatttagcttctacaattaataacatgatgtatcttgata

taccgtttattgcttttactgcaaataacgataattgggtcaagcaagatgaagttatcacattgttatcaaatattcgtagtaatcgatgcaagatatattctttgttaggaagttcgcatgacttgagtgaaaatttagtggtcctgcgcaa

tttttatcaatcggttacgaaagccgctatcgcgatggataatgatcatctggatattgatgttgatattactgaaccgtcatttgaacatttaactattgcgacagtcaatgaacgccgaatgagaattgagattgaaaatcaagcaatt

tctctgtcttaa

*luxE* atgaagggtataaaagagtatgacagcagtgctgccatactttctaatattatcttgaggagtaaaacaggtatgacttcatatgttgataaacaagaaattacagcaagctcagaaattgatgatttgatttttcgagcgatccatta

gtgtggtcttacgacgagcaggaaaaaatcagaaagaaacttgtgcttgatgcatttcgtaatcattataaacattgtcgagaatatcgtcactactgtcaggcacacaaagtagatgacaatattacggaaattgatgacatacct

gtattcccaacatcggtttttaagtttactcgcttattaacttctcaggaaaacgagattgaaagttggtttaccagtagcggcacgaatggtttaaaaagtcaggtggcgcgtgacagattaagtattgagagactcttaggctctgtg

agttatggcatgaaatatgttggtagttggtttgatcatcaaatagaattagtcaatttgggaccagatagatttaatgctcataatatttggtttaaatatgttatgagtttggtggaattgttatatcctacgacatttaccgtaacagaaga

acgaatagattttgttaaaacattgaatagtcttgaacgaataaaaaatcaagggaaagatctttgtcttattggttcgccatactttatttatttactctgccattatatgaaagataaaaaaatctcattttctggagataaaagcctttat

atcataaccggaggcggctggaaaagttacgaaaaagaatctctgaaacgtgatgatttcaatcatcttttatttgatactttcaatctcagtgatattagtcagatccgagatatatttaatcaagttgaactcaacacttgtttctttgag

gatgaaatgcagcgtaaacatgttccgccgtgggtatatgcgcgagcgcttgatcctgaaacgttgaaacctgtacctgatggaacgccggggttgatgagttatatggatgcgtcagcaaccagttatccagcatttattgttacc

gatgatgtcgggataattagcagagaatatggtaagtatcccggcgtgctcgttgaaattttacgtcgcgtcaatacgaggacgcagaaagggtgtgctttaagcttaaccgaagcgtttgatagttga

*asd* atgtgccaggaggagaccggcacatttatacagcacacatctttgcaggaaaaaacgcttatgaaaaatgttggttttatcggctggcgcggtatggtcggctccgttctcatgcaacgcatggttgaagagcgcgacttcgacgc

cattcgccctgtcttcttttctacttctcagcttggtcaggctgcgccgtcttttggcggaaccactggcacacttcaggatgcctttgatctggaggcgctaaaggccctcgatatcattgtgacctgtcagggcggcgattataccaac

gaaatctatccaaagcttcgtgaaagcggatggcaaggttactggattgacgcagcatcatctctgcgcatgaaagatgacgccatcatcattcttgaccccgtcaatcaggacgtcattaccgacggattaaataatggcatca

ggacttttgttggcggtaactgtaccgtaagcctgatgttgatgtcgctgggtggtttattcgccaatgatcttgttgattgggtgtccgttgcaacctaccaggccgcttccggcggtggtgcgcgacatatgcgtgagttattaacccaa

atgggccatctgtatggccatgtggcagatgaactcgcgaacccgtcctctgctattctcgatatcgaacgcaaagtcacaaccttaacccgtagcggtgagctgccggtagataactttggcgtgccgctggcgggtagcctgat

tccgtggatcgacaaacagcttgataacggtcagagccgcgaagagtggaaagggcaggcggaaaccaacaagatcctcaacacatcttccgtaattccggtagatggtttatgtgtgcgtgtcggggcattgcgctgccaca

| | |
|---|---|
| | gccaggcattcactattaaattgaaaaaagatgtgtctattccgaccgtggaagaactgctggctgcgcacaatccgtgggcgaaagttgttccaaacgatcgggaaatcactatgcgtgagctaaccccagctgccgttaccg |
| | gcacgctgaccacgccggtaggccgtctgcgtaagctgaatatgggaccagagttcctgtcagcctttaccgtgggcgaccagctgctgtggggggccgcggagccgctgcgtcggatgcttcgtcaactggcgtaa |
| *coaA* | atgaaaaagcagatcatcagcctgggtgcactggcagtagcaagcagcctgtttacctgggataacaaagcagatgcaattgtgaccaaagattatagcggtaaaagccaggttaatgcaggtagcaaaaacggcaccctg |
| | attgatagtcgttatctgaatagcgcactgtattatctggaagattatattatctatgcaatcggtctgaccaataaatatgaatatggtgataacatctataaagaagcaaaagatcgtctgctggaaaaagttctgcgtgaagatca |
| | gtatctgctggaacgtaaaaagagccagtatgaagattataaacagtggtatgcgaattataaaaaggaaaatccgcgtacagatctgaaaatggccaactttcataaatataacctggaagaactgagcatgaaagaatata |
| | atgaactgcaggatgcactgaaacgtgcactggatgattttcatcgcgaagttaaagatattaaagataaaaacagcgatctgaaaacctttaatgcagcagaagaagataaagcaaccaaagaagtatatgatctggtgtctg |
| | aaatcgatacactggtcgtaagctattatggtgataaagattatggcgaacatgccaaagaactgcgtgcaaaactggatctgatcctgggcgatacggataatcctcataaaattaccaacgaacgtattaaaaaggaaatga |
| | tcgatgatctgaacagcattattgatgattttttatggaaaccaaacagaaccgtccgaaaagcattacaaaatataacccgaccacccataactataaaaccaacagcgataacaaaccgaattttgataaactggtggaag |
| | aaaccaaaaaggcagtcaaagaagcagatgatagctggaaaaagaaaaccgtgaaaaagtatggtgaaaccgaaaccaaaagcccggtagtgaaagaagaaaaaaaagttgaagaaccgcaggcaccgaaagtt |
| | gataatcagcaggaagttaaaaccaccgcaggtaaagcagaagaaaccacccagccggttgcacagccgctggttaaaattccgcagggtaccattaccggtgaaattgttaaaggtccggaatatccgacaatggaaaat |
| | aaaaccgttcagggtgaaattgttcagggtccggattttctgacaatggaacagagcggtccgagcctgagcaataattataccaatccgccgctgaccaatccgattctggaaggtctggaaggtagcagcagcaaactgga |
| | aattaaaccgcagggtaccgaaagcaccctgaaaggtacccagggtgaaagcagcgatattgaagttaaaccgcaggcaaccgaaaccaccgaagcaagccagtatggtccgcgtgttaccaaataa |
| luxI | atgactataatgataaaaaaaatcggatttttggcaattccatcggaggagtataaaggtattctaagtcttcgttatcaagtgtttaagcaaagacttgagtgggacttagttgtagaaaataaccttgaatcagatgagtatgataact |
| | caaatgcagaatatatttatgcttgtgatgatactgaaaatgtaagtggatgctggcgtttattacctacaacaggtgattatatgctgaaaagtgtttttcctgaattgcttggtcaacagagtgctcccaaagatcctaatatagtcgaa |
| | ttaagtcgttttgctgtaggtaaaaatagctcaaagataaataactctgctagtgaaattacaatgaaactatttgaagctatatataaacacgctgttagtcaaggtattacagaatatgtaacagtaacatcaacagcaatagagc |
| | gatttttaaagcgtattaaagttccttgtcatcgtattggagacaaagaaattcatgtattaggtgatactaaatcggttgtattgtctatgcctattaatgaacagtttaaaaaagcagtcttaaattaa |
| luxR | atgatatataacacgcaaaacttgcgacaaacaataggtaaggataaagagatgggtatgaaaaacataaatgccgacgacacatacagaataattaataaaattaaagcttgtagaagcaataatgatattaatcaatgctt |
| | atctgatatgactaaaatggtacattgtgaatattatttactcgcgatcatttatcctcattctatggttaaatctgatatttcaattctagataattaccctaaaaaatggaggcaatattatgatgacgctaatttaataaaatatgatcctat |
| | agtagattattctaactccaatcattcaccaattaattggaatatatttgaaaacaatgctgtaaataaaaaatctccaaatgtaattaaagaagcgaaaacatcaggtcttatcactgggtttagtttccctattcatacggctaacaat |
| | ggcttcggaatgcttagttttgcacattcagaaaaagacaactatatagatagtttatttttacatgcgtgtatgaacataccattaattgttccttctctagttgataattatcgaaaaataaatatagcaaataataaatcaaacaacgat |
| | ttaaccaaaagagaaaaagaatgtttagcgtgggcatgcgaaggaaaaagctcttgggatatttcaaaaatattaggctgcagtgagcgtactgtcactttccatttaaccaatgcgcaaatgaaactcaatacaacaaaccgct |
| | gccaaagtatttctaaagcaattttaacaggagcaattgattgcccatactttaaaaattaa |
| *SAH* | atggactacaaggacgacgatgacaagaaaacccgtatcgtttcttctgttaccaccaccctgctgctgggctctatcctgatgaacccggttgcgaacgcggcggactctgacatcaacatcaaaaccggcaccaccgacat |
| | cggctctaacaccaccgttaaaaccggcgacctggttacctacgacaaagaaaacggcatgcacaaaaaaagttttctactctttcatcgacgacaaaaaccacaacaaaaaaactgctggttatccgtaccaaaggcaccatc |
| | gctggccagtaccgtgtatacagcgaagaaggcgcgaacaaatctggcctggcgtggccgtctgcgttcaaagttcagctgcagctgccggacaacgaagttgcgcagatatctgactactacccgcgtaactctatcgacac |
| | caaagaatacatgtctaccctgacctacggcttcaacggcaacgttaccggcgacgacaccggcaaaatcggcggcctgataggcgcgaacgtaagcatcggccacaccctgaaatacgttcagccggacttcaaaacca |

tcctggaatctccgaccgacaaaaaagttggctggaaagttatcttcaacaacatggttaaccagaactggggcccgtacgaccgtgactcttggaacccggtttacggcaaccagctgttcatgaaaacccgtaacggctctat
gaaagcggcggaaaacttcctggacccgaacaaagcgtcttctctgctgtcttctggcttctctccggacttcgcgaccgttatcactatggaccgtaaagcgtctaaacagcagaccaacatcgacgttatctacgaacgtgttcg
tgacgactaccagctgcactggacctctaccaactggaaaggcaccaacaccaaagacaaatggaccgaccgttcttctgaacgttacaaaatcgactgggaaaaagaagaaatgaccaactaa
